# Natural variations of cardiac performance in *Drosophila* identify a central function for *Pdp1*/*dHLF* in cardiac aging

**DOI:** 10.1101/2024.09.30.615759

**Authors:** Katell Audouin, Saswati Saha, Laurence Röder, Sallouha Krifa, Nathalie Arquier, Laurent Perrin

**Affiliations:** Aix Marseille Univ, INSERM, TAGC, Turing Center for Living systems, Marseille, France; Argenx France SAS, 24, rue du Gouverneur Général Eboué, Issy les Moulineaux, France; Strasbourg University, CNRS UPR9022, Strasbourg, France; CNRS, Marseille, France

**Author notes:** Equal contribution (Katell Audouin, Saswati Saha). Co-last authors (Nathalie Arquier, Laurent Perrin).

**Keywords:** Association study, cardiac performance, aging, *Pdp1*, mitochondria, *Drosophila*

## Abstract

The identification of genetic factors influencing cardiac senescence in natural populations is central to our understanding of cardiac aging and to identify the etiology of associated cardiac disorders in human populations. However, the genetic underpinning of complex traits in human is almost impossible, due to the infeasibility to control genetic background and gene-environment interactions. *Drosophila* has striking similarities in cardiac aging with humans, highlighting the conserved nature of cardiac aging for organisms with a heart. Leveraging on a large collection of inbred lines from the Drosophila Genetic Reference Panel (DGRP), we provide an accurate analysis of cardiac senescence in a natural population of flies. This permitted the discovery of an unprecedented number of variants and associated genes significantly associated to the natural variation of cardiac aging. We focused on the function of the PAR-domain bZIP transcription factor Pdp1 for which several variants were found associated with natural variation of the aging of multiple cardiac functional traits. We demonstrated that *Pdp1* cell autonomously plays a central role in cardiac senescence and might do so by regulating mitochondria homeostasis. Overall, our work provides a unique resource regarding the genetics of cardiac aging in a natural population.

## Introduction

As for most traits in human populations, cardiac aging and common associated diseases and disorders have a complex genetic basis. Their manifestations depend on multiple, often interacting, segregating genes that also interact with environmental factors. Still, description and functional testing of the genetic architecture underlying cardiovascular aging is of fundamental importance, as it will impact our understanding of the biological basis of associated diseases. This however requires not only the identification of a detailed part list of genes affecting these traits, but also of the actual causal variants, their allele frequencies, molecular effects and individual as well as collective effects on organismal phenotypes.

The identification of loci at which alleles with subtle effects segregate in human populations through Genome Wide Association Studies (GWAS) has been performed for many complex traits including cardiovascular diseases. This led to the identification of large collections of loci potentially linked to cardiac function and disorders^1–4^. However, due to large blocks of linkage disequilibrium in the human genome, the identification of causal variants and genes is challenging. Furthermore, the genetic underpinning of complex traits in human populations is complicated by gene-gene interactions (epistasis) which may result in failure to reproduce associations in different populations with different genetic backgrounds, and by genotype-environment interactions which may result in lack of reproducibility between populations, when associations that are significant in one population in one environment cannot be resolved in another population under different environmental conditions. Altogether this makes the understanding of genetic architecture of complex traits a daunting challenge.

There is therefore a compelling need for comparative studies in model organisms. The genetically tractable *Drosophila* model system has proven to be a potent model in elucidating conserved genetic networks of heart development and of adult heart function and aging^5–10^. Human and fly hearts share many homologous and orthologous genes and gene products, including sarcomeric proteins and ion channels. Heart function can easily be manipulated *in vivo* and the effects of these manipulations are not immediately lethal (O_2_ is delivered by a separate tracheal system) allowing quantitative analysis of malfunctioning hearts^11,12^. A number of cardiac aging studies in *Drosophila* have revealed striking similarities in cardiac aging with vertebrates/humans, both in terms of heart physiology and transcriptional changes^10,13–16^. This highlights the conserved nature of cardiac aging for organisms with a heart.

We recently investigated the natural variations of cardiac performances among a natural population of young (1 week) adult female flies. We used the Drosophila Genetic Reference Panel (DGRP), a community resource constituted of a panel of inbred lines that have been created by mating full siblings of wild-caught iso-female lines for 20 generations^17^. The use of inbred lines allows for assessing the effect of genetic variations on distinct but constant genetic backgrounds and teasing out the effect of genotype from environmental effects. We have used an established *in situ* heart preparation protocol that permits precise quantification of heart function in the fly^11^. We demonstrated substantial among-lines variations of cardiac performances and identified genetic variants associated with the cardiac traits that were extensively validated *in vivo*. This pointed out a central role of gene regulatory network deviations in the genetic architecture of these complex traits. Importantly, we documented correlations between genes associated with cardiac phenotypes in both flies and humans, which supports a conserved genetic architecture regulating adult cardiac function from arthropods to mammals^18^.

Here we report an in-depth characterization of the aging of cardiac performance traits in flies, starting from a large panel of individuals in the DGRP, from 193 inbred lines analyzed both in young and aging females. Using statistical models based on individual phenotypes, the genome-wide association studies identified a large collection of variants that remained significant genome-wide after Bonferroni correction for multiple testing. Importantly, we found that a significant proportion of human orthologs of identified genes were identified in GWAS studies for cardiac disorders, further highlighting the conserved nature of cardiac functioning. We focused on the role of transcription factors (TFs) in the genetic architecture of natural variation of cardiac aging. We identified a central function for *PAR-domain protein 1* (*Pdp1*) gene - encoding a bZIP TF orthologous to human HLF and TEF TFs - in the control of senescence of both cardiac contractility and rhythmicity. We also provide several lines of evidence indicating that *Pdp1* may affect cardiac aging by regulating mitochondria number and physiology.

## Results

### Screening the DGRP for aging of heart performances

We previously reported the analysis of the genetic architecture of natural variations of cardiac performance traits in young adult females in the DGRP. Heart parameters were measured in 1-week-old females for 167 DGRP lines^18^ using high-speed video recording of the heart beating on semi-intact preparations of individual flies^19,20^. Here, we wanted to identify the genetic variants associated with the age-dependent modifications of cardiac performance traits. 4 weeks old female flies of the same DGRP lines were therefore analyzed for cardiac performances using the same setup (12 females per lines). The high mortality rate we -and others^21^-observed in a significant proportion of DGRP lines from 30 days onward for standard laboratory temperature of 25°C prevented the analysis at older ages. A batch of 26 additional lines were analyzed at both ages, bringing the total number of lines analyzed to 193, representing a total of more than 4000 individual flies (2297 flies at 1 week, 2084 flies at 4 weeks, see Supplemental Dataset 1).

High-speed videos were analyzed using the Semi automated Heartbeat Analysis software (SOHA, http://www.sohasoftware.com/) which allows quantification of several cardiac functional parameters (Figure 1A) allowing 8 different traits to be measured on each individual. We analyzed cardiac traits related to rhythmicity – systolic intervals (SI, time elapsed between the beginning and the end of the contraction), diastolic intervals (DI, time elapsed between two contractions), heart period (HP, the duration of a total cycle (HP= DI + SI)) and arrhythmia index, to evaluate the variability of the cardiac rhythm (AI, std-dev(HP)/ mean (HP)). We also analyzed cardiac traits related to contractility – the diameters of the heart in diastole (end diastolic diameter, EDD), in systole (end systolic diameters, ESD), and the fractional shortening which measures contraction efficiency (FS = (EDD-ESD)/EDD). In addition, cardiac output - which measures the volumetric flow rate of the heart (CO = SV/HP, where SV is the stroke volume (volume of hemolymph pumped by the heart per beat)) - was also analyzed.

**Figure 1.**
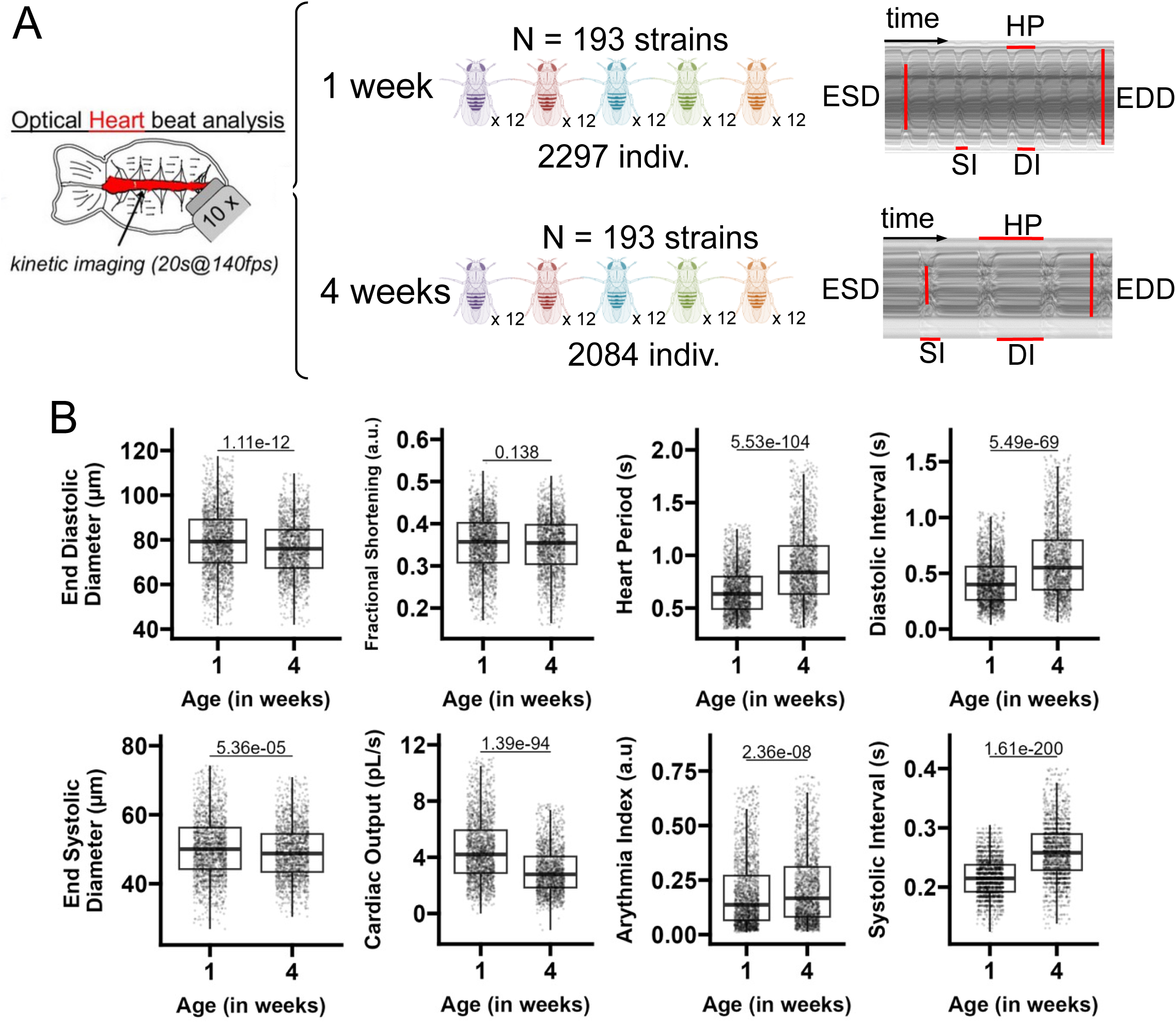
Cardiac aging in the DGRP. (A) Cardiac performance traits were analyzed on 1 week and 4 weeks old females flies from 193 DGRP lines on semi-intact preparations of *ex-vivo* isolated hearts. Up to 12 individuals per DGRP line were analyzed. Total number of individuals examined are indicated. Right: Representative M-modes at 1 week and 4 weeks. DI=Diastolic Interval; SI=Systolic Interval; HP=Heart Period; EDD=End Diastolic Diameter; ESD=End Systolic Diameter. Fractional Shortening (FS=EDD - ESD/EDD), Arrythmia index (AI=Std Dev(HP)/HP) and Cardiac output (CO=(π(EDD²/2) - π(ESD²/2)) /HP) were also measured. (B) Boxplot of all individual fly phenotypes compared between 1 and 4 weeks in the whole DGRP population analyzed. Wilcoxon rank sum test was applied to test the significance between 1- and 4-weeks values.

### Phenotypic description of heart aging in the DGRP

The 8 cardiac traits were compared between young (1 week) and aged (4 weeks) flies after performing quality control checks on the phenotype data (see Materials and methods). Strong aging trends were observed across the different traits (see Figure 1B). Given the size of the sample analyzed, our phenotypic description of drosophila cardiac functional senescence in a natural population is unprecedented. Except for fractional shortening, which did not significantly change from 1 to 4 weeks in the DGRP population, all other cardiac performance traits displayed significant age-related modifications. In old flies, heartbeats were lengthened at all phases of the cardiac cycle (HP, DI, and SI were longer at 4 weeks compared to one week) and the arrhythmias index increased, slightly but significantly. In addition, we noticed a constriction of the heart both in diastole and in systole, with a reduction of EDD and ESD with age. Accordingly, cardiac output – which is a function of both heart rate and cardiac volumes-was strongly reduced at week 4. Figure 2A and Supplemental Figure 1A illustrate how, for each DGRP line, average cardiac trait values evolve from 1 to 4 weeks. Even in the case of FS, which at the level of the whole population appears not to be affected by age (Figure 1B), DGRP lines show great variability in trends with age, suggesting a major influence of genetic factors that influenced the evolution of FS with age. Overall, the phenotypes were poorly correlated with each other, with the exception of HP with DI, EDD with ESD and, to a lesser extent, HP with SI (Figure 2B).

**Figure 2.**
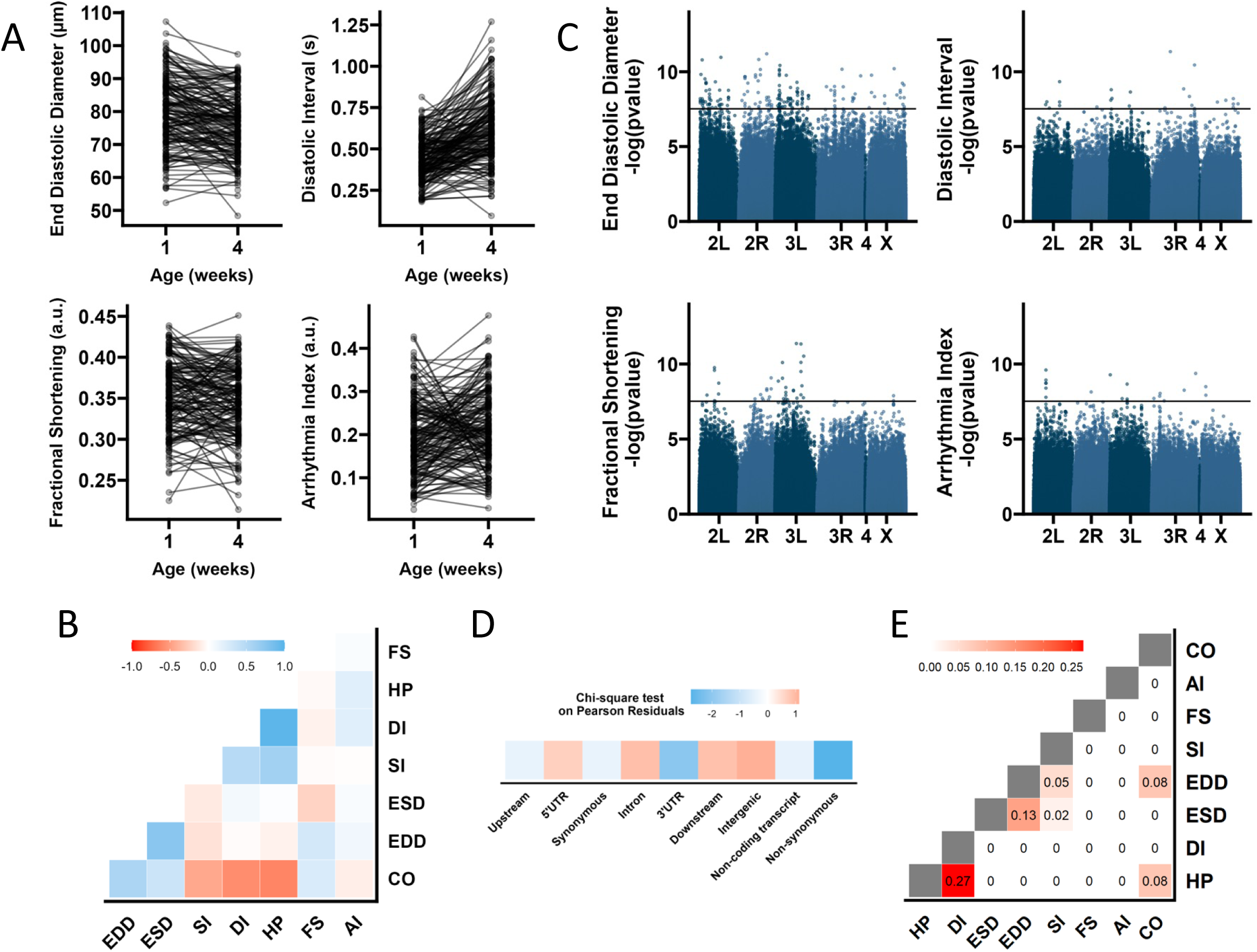
Genome-Wide associations studies (GWAS) for aging of cardiac performance in the DGRP. (A) Reaction norm plot of phenotypic mean for all DGRP lines at 1 and 4 weeks for EDD, FS, DI and AI phenotypes. The mean phenotypic values for each line at each age are connected by a line, allowing to monitor the trend differences among DGRP lines. (B) Overall correlations between senescence of cardiac traits in the DGRP. Spearman correlations were calculated based on the difference between the average of line means at each age for each cardiac trait. (C) Manhattan plot showing the results of GWAS performed on cardiac traits modifications with age. The genome wide significance threshold (3e-8) is indicated by a black line. (D) Pearson residuals of chi-square tests from the comparison of randomly selected SNP in the DGRP with variants associated with cardiac traits. According to Variant Effect Predictor (VEP) and a custom pipeline of annotation priorization, SNPs are attributed to different localization categories. SNPs are attributed to genes if they are within the gene transcription unit (5’ and 3’ UTR, synonymous and non-synonymous coding, introns) or within 5 kb from transcription start site (TSS) and transcription end site (TES) (5 kb upstream, 5 kb downstream). Intergenic: SNPs not attributed to genes (>5 kb from TSS and TES). (E) Overlap coefficient of gene sets associated with the different cardiac traits analyzed.

To assess the influence of genetic diversity on the evolution of cardiac traits we have computed the broad sense heritability (H2) (see Materials and methods) for each phenotype. The broad sense heritability ranged from 0.165 (AI) to 0.399 (SI) (Table 1).

**Table 1:** Summary statistics over all DGRP genotypes assayed. Number of lines available for GWAS in each phenotype and at each age after QC check (see Materials and methods). Broad sense heritability (H2) of evolution of traits means between 1 and 4 weeks suggested heritability of corresponding traits.

| Cardiac traits | Nb lines 1w | Nb line 4w | Nb line 1w and 4w | $H^2$ (1 to 4 w) |
| --- | --- | --- | --- | --- |
| Arrhythmia Index | 181 | 178 | 166 | 0.165 |
| Cardiac Output | 178 | 168 | 155 | 0.271 |
| End Diastolic Diameter | 183 | 170 | 161 | 0.272 |
| End Systolic Diameter | 178 | 169 | 158 | 0.229 |
| Diastolic Interval | 186 | 178 | 172 | 0.258 |
| Fractional Shortening | 183 | 167 | 159 | 0.209 |
| Heart Period | 186 | 178 | 172 | 0.279 |
| Systolic Interval | 190 | 174 | 171 | 0.399 |

### Performing GWAS for natural variations of aging of cardiac performance

To identify the potential candidate variants associated with the aging of cardiac traits, we performed a single marker GWAS using a linear mixed model on the DGRP phenotypes at 1 week and 4 weeks, accounting for the effects of Wolbachia infection and common polymorphism inversions (Figure 2C and Supplemental Figure 1; see Materials and methods). GWAS was performed separately for all eight cardiac traits and SNPs/variants were ranked based on their p-values.

SNPs were selected after a stringent statistical correction for false positives based on Bonferroni correction, derived by dividing 1,635,932 SNPs into 0.05, giving a genome-wise significance threshold of 3×10e-8 (Supplemental Table 1, Figure 2C, and Supplemental Figure 1; see Materials and Methods and Supplemental Materials and Methods). The number of SNPs selected using the above threshold varied from 12 (CO) to 178 (EDD) and, overall, 380 unique variants were identified (see Table 2). Note that, in most DGRP studies, GWAS are performed on the mean of the phenotypes for DGRP lines, thereby restricting the power of the studies. Here, the linear mixed model (LMM) was applied on individual cardiac performance traits of a large population of flies (> 8 individuals per line and per age, see Materials and Methods). The QQ plots of the GWAS p-values across all phenotypes showed some inflation (Supplemental Figure 3). However, it is well known that a certain degree of inflation is commonly observed in interaction GWAS analyses^22,23^ particularly due to reduced power and model complexity. To investigate the potential source of this inflation, we conducted additional analyses described in detail in the Supplemental Material and Methods. These analyses indicate that the observed inflation is unlikely to arise from misspecification of the interaction model. Instead, when evaluated against an empirical null distribution tailored to our data structure, the leading interaction signals remain robust and largely preserved. Taken together, these results provide strong evidence that the leading interaction hits identified in our GWAS are unlikely to represent false positives.

**Table 2.**
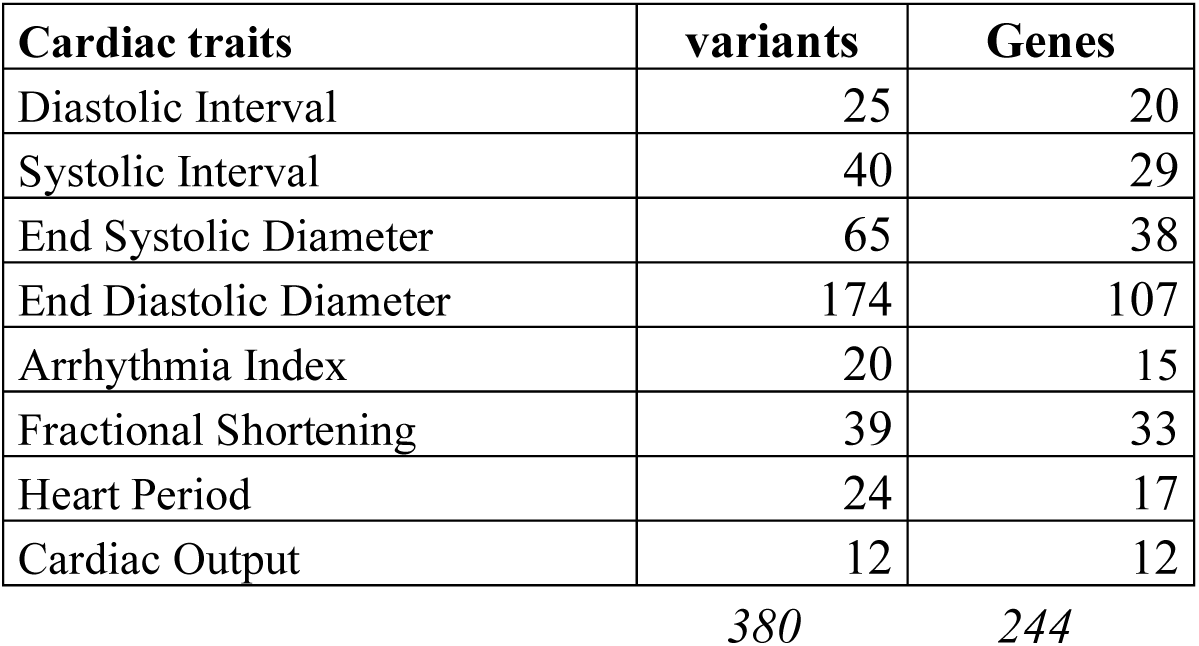
Number of variants and genes identified by GWAS for the aging of cardiac performance trait. Total number of (non-redundant) variants and genes identified across all traits are indicated below.

Identified variants were mapped onto the dm6 genome in order to identify associated genes (Supplemental Table 1). Compared to random expectation, variants were depleted for non-synonymous coding alleles and for mutations affecting genes 3’ UTR (Figure 2D), suggesting that most deleterious mutations affecting cardiac aging were counter selected in the population. Selected variants mapped into 12 (CO) to 107 (EDD) genes (Table 2), identifying a total of 244 unique genes across the 8 cardiac traits analyzed. GWAS for senescence of HP and DI identified partially overlapping gene sets (Figure 2E), and the same was true, but to a lesser extent, for GWAS on ESD and EDD. This is in line with the correlations identified among corresponding phenotypes (Figure 2B).

### Functional annotations of associated genes

Remarkably, GWAS for aging of ESD identified a variant within the *vrille* (*vri*) gene (Supplemental Table 1), encoding a conserved bZIP transcription factor that we previously identified as a master regulator of cardiac aging^24^. In addition, the set of genes associated to ESD also included the *Catalase* (*Cat*) gene encoding a reactive oxygen species-scavenging enzyme, whose activity was demonstrated to prevent cardiac senescence^24^. The recovery of variants for both *vri* and *cat* in GWAS for cardiac aging represents strong arguments in favor of the accuracy of our association analyses.

In an attempt to gain insights into the biological processes affected by natural variation of cardiac senescence, we searched for Gene Ontologies annotations overrepresented within the set of genes identified across all GWAS (https://biit.cs.ut.ee/gplink/l/0YrfbOT9RO). Overall, only a few functional annotations were identified as overrepresented among GWAS associated genes – mainly related to development and morphogenesis. Enrichment analyses therefore did not allow the deciphering of enriched molecular and/or cellular processes affected in the genetic architecture of cardiac senescence. Strikingly however, when compared to the whole set of genes in the fly genome, GWAS associated genes are enriched for genes that have an ortholog in the human genome (FC 1,16, Table 3) indicating that, overall, genes associated to natural variations of cardiac aging have evolutionarily conserved functions. In addition, the set of drosophila genes identified here, throughout GWAS for natural variations of cardiac aging, is enriched for genes whose human orthologs have been found associated to cardiac disorders (FC 1,5) and to CAD (Coronary Artery Disfunction; FC 1,69), suggesting that cardiac aging in flies and cardiac pathologies in humans share an overlapping genetic basis.

**Table 3:** Conserved genes associated with natural variations of aging of cardiac traits from flies to humans. Enrichment analyses for genes conserved in human and for genes whose human ortholog is associated with either coronary artery diseases (CAD) or cardiac disorders. Second row (human orthologs): Number of fly genes that display a human ortholog according to DIOPT (https://www.flyrnai.org/diopt; high and moderate rank, Supplemental Table 2). Third and fourth rows: Number of genes whose human ortholog (high and moderate rank) has been associated with CAD or cardiac disorders by genome-wide associations studies (GWAS) in human populations^18^. Fold change (FC): Ratio between expected (based on successes observed on fly genome) and observed number of successes in respective GWAS gene sets. pval: hypergeometric p-value.

|  | Fly genome | GWAS Aging | Fold Change | pvalue |
| --- | --- | --- | --- | --- |
| Fly genes | 17895 | 244 |  |  |
| Human orthologs | 10647 | 168 | 1.16 | 0.001 |
| Coronary Artery Diseases | 451 | 12 | 1.69 | 0.027 |
| Cardiac Disorders | 1225 | 29 | 1.50 | 0.007 |

The majority of the variants identified across all GWAS are located in introns, downstream and upstream of the candidate genes, or in intergenic regions (Supplemental Table 1), suggesting that they affect cardiac senescence via modulation of gene expression. This may represent a common feature of the genetics underpinning of natural variation of complex traits. As a matter of facts, we already established that natural variation of cardiac performance in young flies primarily affect gene regulatory networks and identified a number of transcription factors (TFs) involved^18^.

Table 4 lists all the TFs identified in the present study. Among all identified TFs, a few were retrieved in different GWAS and therefore associated to the natural variation of several cardiac aging traits. This in particular was the case for *PAR-domain protein 1* (*Pdp1*), an ortholog of mammalian *HLF*, *DBP* and *VBP/TEF* bZip transcription factors. Variants within the *Pdp1* gene were associated to variations of aging of ESD, EDD and SI, suggesting its involvement in the natural variations in the aging of both rhythmicity and contractility.

**Table 4:**
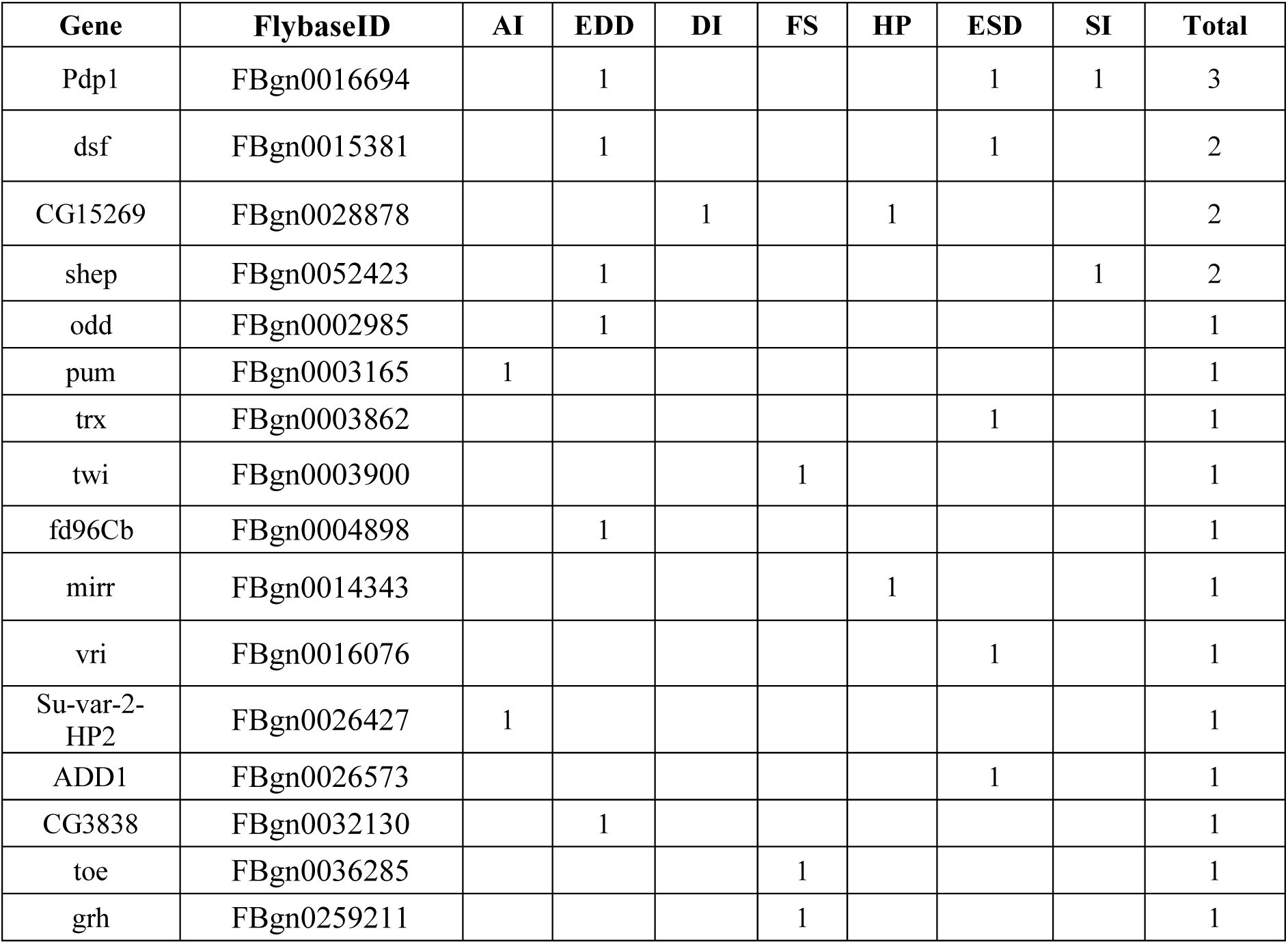
List of genes encoding transcription factor identified across GWAS for cardiac aging. Traits for which variants in genes were associated ((p<3e-8) are indicated. Genes are ranked according to the number of traits for which they were identified, across all 8 GWAS (column ‘total’).

### Pdp1, a bZip transcription factor with multiple effects on cardiac senescence

The *Pdp1* gene encodes multiple protein isoforms and is involved in a variety of biological processes. It regulates muscle genes expression during differentiation and was shown to control that of *Tropomyosin I*, possibly throughout interaction with Mef2^25^. It also regulates cellular lipid contents in fat body, similarly to mammalian HLF^26^. *Pdp1* is also involved in organismal growth control and its expression was shown to be sensitive to nutritional conditions^27^. In addition, it was characterized as an interactor of *vri* in clock neurons, forming together with *clock*, a second feedback loop in the drosophila circadian clock^28^. There, Pdp1 ε isoform and Vri display antagonistic activity, being respectively direct activator and repressor of the *clock* gene, by direct binding on the same DNA motif^28^. It was therefore striking to identify both *vri* and *Pdp1* in GWAS for natural variations of cardiac aging. As mentioned above, *vri* impact on cardiac aging has been characterized previously and its loss of function significantly reduces age-related cardiac dysfunctions^24^. Hence, we investigated *Pdp1* function in the heart during aging, taking advantage of a heart-specific inducible driver^24^. This allowed us to perform adult specific gene knock-down (KD) and over-expression (OE), independently of developmental effects. As shown in Figure 3 and Supplementary Figure 4, *Pdp1* KD markedly accelerated senescence of AI - whose increase throughout aging was amplified- and also enhanced the reduction of both systolic and diastolic diameters with age. On the contrary, *Pdp1* KD appeared to delay the effects of age on traits related to cardiac rhythm. We indeed observed a mild - but significant - reduction of SI increase with age, and a reduced DI and HP with age, while both parameters display an increased trend in controls. Similar trends, although less pronounced, were observed using an independent RNAi construct (Supplemental Figure 5). Remarkably, the effects of *Pdp1* OE were opposite to those observed following its KD (Figure 3B). It markedly prevented AI increase from 1 to 7 weeks. Also, although the aging trend was not significant, *Pdp1* OE led to cardiac hypertrophy with age, more noticeable in diastole. Conversely, HP, DI, and SI increase with age were amplified. Overall, *Pdp1* contribution to cardiac aging appears to be complex, with beneficial (HP, DI and SI) or detrimental (ESD, DD, AI) effects, depending on the traits considered.

**Figure 3.**
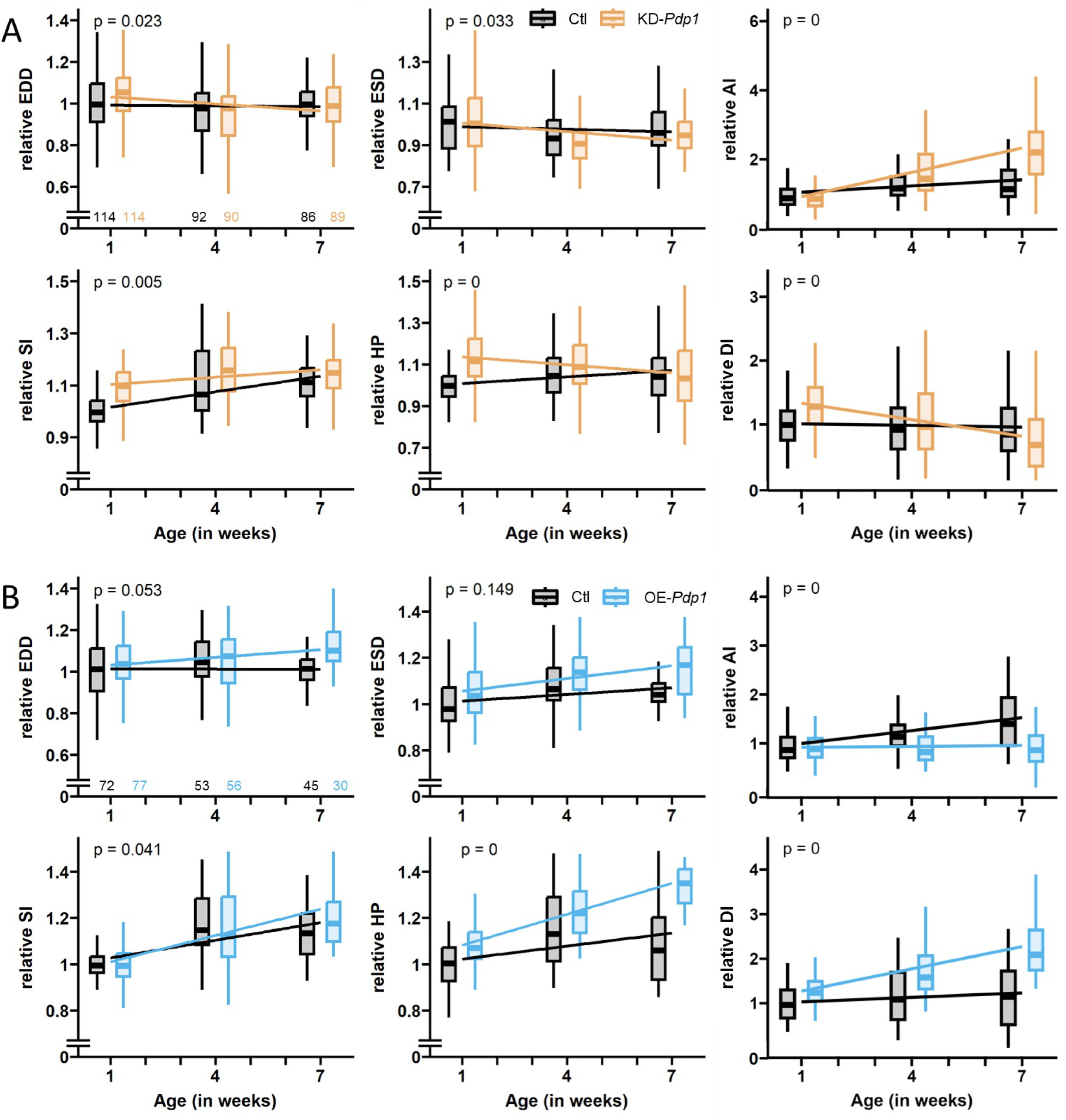
Heart-specific conditional knock-down (KD) or overexpression (OE) of *Pdp1* modulate cardiac senescence. Cardiac performance was analyzed on intact anesthetized females flies at indicated ages. Number of individuals analysed are indicated on the first plot for each genotype. Individual values were normalized to the mean of the control condition at 1 week. Linear model was applied to analyze the statistical difference between aging trends of control and either *Pdp1* knock-down (A) or over-expression (B). (A) Effect of *Pdp1* RNAi mediated gene knockdown (KD). w/w; tdtk/UAS>dsRNA(*Pdp1*)^VDRC-KK^; Hand>GeneSwitch-Gal4 /+ flies fed (KD-*Pdp1*) or not (Ctl) with 100µg/ml of RU486 from adult eclosion onward were imaged at 1 week, 4 weeks and 7 weeks. (B) Effect of *Pdp1* over expression (OE). w/w; tdtk/UAS>*Pdp1*; Hand>GeneSwitch-Gal4 /+ flies fed (OE-*Pdp1*) or not (Ctl) with 100µg/ml of RU486.

### *Pdp1* may affect heart function through the regulation of mitochondria homeostasis

We previously reported that the set of potential Vri cardiac targets during the aging process was strongly enriched in genes encoding mitochondrial proteins^24^, suggesting that *vri* affects cardiac senescence by impinging on mitochondria function. Several variants located in nuclear gene encoding proteins involved in mitochondria function and physiology were retrieved in our GWAS (Table 5), supporting the implication of mitochondria (dys)function in cardiac aging. In addition, in an attempt to decipher the gene regulatory network(s) that may be affected by natural variation of cardiac aging, we used the i-cistarget method, which takes advantage of a comprehensive library of TF-binding motifs and of ChIP-seq experiments together with ranking statistics to predict potential regulatory TFs among set of genes^29^. Here, we aimed to identify potential regulators of gene sets that have GWAS identified variants associated to the natural variation of aging of cardiac traits. Remarkably, when a gene set associated to natural variations of ESD were used, Vri and Pdp1 were predicted as potential regulators by i-cisTarget (https://gbiomed.kuleuven.be/apps/lcb/i-cisTarget/reports/e37e86a59b1774e6c4512368f40f7d4b40d42c81/report.html). In addition, i-cisTarget predicted Vri and Pdp1 target genes included themselves together with *milt* and *larp*, that respectively regulate mitochondria transport^30–32^ and mitochondrial protein translation-thus impinging on mitochondrial DNA (mtDNA) replication^33^.

**Table 5:** List of genes encoding proteins involved in mitochondria physiology identified across GWAS carried out for cardiac aging. Traits for which variants in these genes were associated (p<3e-8) are indicated.

| Gene | FlybaseID | EDD | DI | FS | HP | ESD | SI |
| --- | --- | --- | --- | --- | --- | --- | --- |
| milt | FBgn0262872 |  |  |  |  | 1 |  |
| larp | FBgn0261618 |  |  |  |  | 1 |  |
| MCU | FBgn0042185 | 1 |  |  |  | 1 |  |
| MICU1 | FBgn0039029 | 1 | 1 |  | 1 |  |  |
| CG12464 | FBgn0033861 |  |  |  |  |  | 1 |
| mRpL27 | FBgn0053002 |  |  | 1 |  |  |  |
| Atg1 | FBgn0260945 | 1 |  |  |  |  |  |
| CG9399 | FBgn0037715 | 1 |  |  |  |  |  |
| Vsp13D | FBgn0052113 | 1 |  |  |  |  |  |
| pmi | FBgn0044419 | 1 |  |  |  |  |  |
| Cat | FBgn0000261 | 1 |  |  |  |  |  |

Taken together, these observations suggested that, similar to its paralogous gene *vri*, *Pdp1* may affect cardiac function and aging at least in part by impinging on mitochondria function. To test this hypothesis, we tested whether *Pdp1* interacts with *MCU* (Mitochondrial Calcium Uniporter) which encodes the pore-forming subunit of the mitochondrial calcium uniporter channel (mtCU), a multi-protein complex that spans the inner mitochondrial membrane and is involved in the rapid uptake of calcium into the mitochondrial matrix^34–36^. Variants in *MCU* were associated to natural variations of aging of multiple cardiac traits (Table 5). In addition, a variant in *CG4704*, encoding the fly MICU1 ortholog^37^ – one of the accessory regulatory proteins of the mtCU^38^ – was also retrieved in our GWAS (Table 5). The identification of multiple variants in different mtCU components strongly suggested that calcium regulation of mitochondrial physiology might play a key role during cardiac aging. As shown in Figure 4A-B and Supplemental Figure 6C-D, we found true genetic interactions between *MCU^1^* and *Df(3L)BSC631^Pdp1^*for several cardiac functional parameters. Indeed, while flies heterozygous for loss of function mutations for either *MCU* (*MCU^1^*) or *Pdp1* (*Df(3L)BSC631^Pdp1^*) displayed no or weak effects on cardiac traits, *MCU^1^*; *Df(3L)BSC631^Pdp1^* double heterozygous flies displayed phenotypes that were significantly different from both control flies and flies carrying single heterozygous alleles for several cardiac traits (HP and SI), or from control flies alone (MAD-HP and MAD-DI – measuring HP and DI variations illustrating cardiac arrhythmias). These results are strong support for an involvement of *Pdp1* in regulating mitochondria function in the heart.

**Figure 4.**
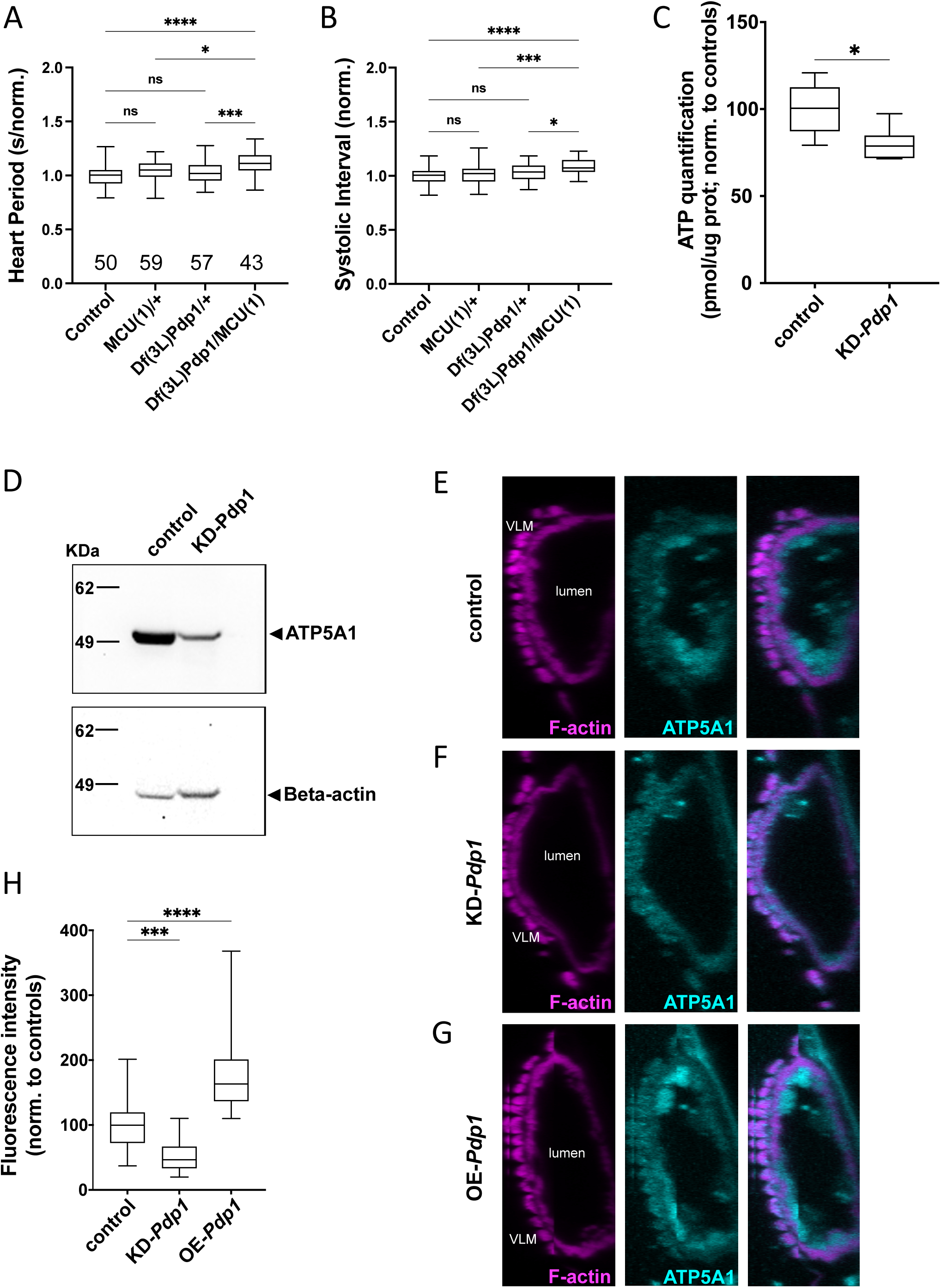
*Pdp1* loss of function genetically interacts with *MCU* and affects mitochondria function. (A-B) Cardiac performance traits analyzed on adult female flies at 1 week (A: Heart Period (HP) and B: Systolic interval (SI)). <u>Control</u>: w/w; tdtk/+; *MCU^1^*: w/w;tdtk/+;*MCU^1^*/+; <u>Df(3L)*Pdp1*</u>: w/w;tdtk/+;Df(3L)BSC631^Pdp1^/+ and MCU^1^/ Df(3L)*Pdp1*: w/w;tdtk/+; MCU^1^/Df(3L)BSC631^Pdp1^. Individual values were normalized to the mean of the control condition. Wilcoxon test was applied to calculate statistical differences between conditions. (C) ATP quantification from isolated thoraces (control: Mef2>RNAi-control, n=46 thorax in 5 independent samples; KD-*Pdp1*: w/w;Mef2-Gal4/ UAS>dsRNA(Pdp1)^VDRC-KK^, n=51 thorax in 6 independent samples). ATP concentration was adjusted to protein content and normalized to the mean of the control condition. Unpaired t test with Welch’s correction was applied. (D) ATP5A1 detection by Western Blot on thorax lysates (top) and Beta-actin normalization (bottom) after membrane stripping. (Control: Mef2>RNAi-control; KD-*Pdp1*: w/w;Mef2-Gal4/ UAS>dsRNA(Pdp1)^VDRC-KK^). 3 independent experiments (10 thoraces per condition) were performed. (E-G) Confocal imaging of hearts stained for ATP5A1 (green) and filamentous actin (F-actin, red) of control (E: w/w;Hand-GAL4/+); heart specific *Pdp1* knock down (F: w/w;Hand-GAL4/ UAS>dsRNA(Pdp1)^VDRC-KK^) and heart specific *Pdp1* overexpression (G: w/w;Hand-GAL4/+;UAS-Pdp1 /+). Individuals were kept at 18°C throughout development, shifted at 25°C at adulthood and imaged at 1 week. Reconstructed orthogonal view of the heart at segment A3 are shown. VLM=Ventral Longitudinal Muscles. (H) Fluorescence quantification of ATP5A1 signals from segments A2 and A3 calculated using the average gray intensity of 200*200px squares. 3-4 cardiomyocytes in 7-11 hearts were analyzed. One-way anova with Dunnett’s correction for multiple testing was applied to calculate statistical difference between control and *Pdp1* conditions. * (p<0,05) / ** (p<0,01) / *** (p<0,001) / **** (p<0,0001)

Since mitochondria’s main function is to supply cells with energy through respiration-associated ATP synthesis, we next tested whether modulating *Pdp1* function in muscle cells was associated to ATP content modifications. We used the muscle specific Mef2 driver to knockdown *Pdp1* function. We first observed that adult flies with reduced *Pdp1* function in their muscles were unable to fly and exhibited severely limited and slow movements compared to controls, suggesting a reduction of their energy capacity (Supplemental Video 1). ATP content was measured on thoraces, which are mainly composed of indirect flight muscles (IFM). As shown in Figure 4C, this resulted in a significant reduction of ATP content compared to sibling controls, thus suggesting that *Pdp1* function indeed affected cell’s ATP synthesis. In agreement, this was associated to a reduction of mitochondrial ATP synthase content, since *Pdp1* knock down was also associated to a strong reduction of ATP5A1 as shown on western blot performed on thoraces of Mef2>*Pdp1* KD adult flies (Figure 4D, Supplemental Figure 6F). In these experiments, we observed a lethality associated to Mef2 driven *Pdp1* overexpression, thus preventing to test the effect of *Pdp1* gain of function on ATP production. We then analysed mitochondria networks in cardiomyocytes from 1week adult flies with heart specific knock down or over-expression of *Pdp1*. As shown in transversal (Figure 4E-F) and longitudinal (Supplemental Figure 6G-H) sections of hearts stained for both ATP5A1 and F-actin, *Pdp1* KD was associated to a reduction of mitochondria network in cardiac cells, where these organelles appeared restricted to the apical part of the cardiomyocytes, while they formed a much larger network in control cardiomyocytes. In addition, the fluorescent signal from ATP5A1 staining was significantly reduced in *Pdp1* KD cardiomyocytes (Figure 4H). On the contrary, *Pdp1* OE resulted in a strong and significant increase of ATP5A1 staining in the heart (Figure 4E,G; Supplemental Figure 6G,I) thus confirming that *Pdp1* function impinges on mitochondria content in this organ.

## Discussion

In this study, leveraging on a large collection of inbred lines from the DGRP and on the phenotypic characterization of over 4000 individuals, we provide an accurate analysis of cardiac senescence in a natural population of flies. This strategy permitted the discovery of an unprecedented number of variants and associated genes significantly associated to the natural variation of these complex traits. Despite the high number of variants confidently associated to natural variations of cardiac aging, we failed to identify enriched annotations among the corresponding genes. This suggests that the deterioration of cardiac function with age is the result of complex interplays between a range of molecular and cellular processes. We however characterized an enrichment for genes whose orthologs are associated to cardiac disorders in humans, in agreement with the known conservation of cardiomyocytes structure, function and aging^39–41^. Our GWAS results therefore represent a unique resource regarding the genetics of cardiac aging in a natural population.

The genetic architecture of senescence of cardiac traits appears dominated by variants located in non-coding regions (introns, intergenic regions) that most probably affect gene expression regulation. To gain insight into the gene regulatory networks affected by natural variation of cardiac aging, we focused on natural variations that affected *Pdp1*, encoding a bZIP transcription factor paralogous to *vri* which we previously demonstrated to regulate cardiac aging. Importantly, variants within *Pdp1* were independently associated with the senescence of EDD, ESD and SI and both *vri* and *Pdp1* were themselves predicted as transcriptional regulators of a set of genes identified in GWAS for natural variations of aging of ESD. In agreement with our GWAS results, heart specific knock down or overexpression of *Pdp1* profoundly affected the modification of several cardiac traits with age. In the case of traits related to rhythm (HP, DI, SI), *Pdp1* KD prevented the effects of age on these cardiac traits, what supports a function of *Pdp1* in promoting cardiac aging. Conversely, with respect to arrhythmia index and systolic and diastolic diameters, *Pdp1* KD accelerated heart senescence, indicating that *Pdp1* function rather prevented cardiac aging of these specific traits.

Hence *Pdp1* function during cardiac aging appears complex. Opposite contributions to cardiac aging may be supported by distinct isoforms, since the *Pdp1* locus encodes at least 6 isoforms^42^. Further characterization of isoform specific contributions will be necessary to disentangle the contribution of *Pdp1* to cardiac aging.

Here, we present strong arguments that *Pdp1* effects on the modifications of cardiac performances with age may be mediated, at least in part, by its impact on mitochondria physiology. Indeed, *Pdp1*, together with *vri*, was predicted to regulate genes implicated in the control of mitochondria activity and we characterized a genetic interaction between *Pdp1* and MCU, a key component of calcium mitochondria homeostasis. In addition, we showed that reducing *Pdp1* activity in muscles resulted in lethargic flies and this was associated with reduced ATP level and ATP synthase contents. Finally, in cardiomyocytes, *Pdp1* activity was shown to be associated with the size of mitochondria network and significantly correlated with ATP synthase content.

Given the central role of mitochondria in energy metabolism and cell survival, the heart is highly susceptible to perturbations of mitochondria homeostasis. Mitochondrial influence on age related cardiac dysfunctions is therefore largely recognized^43,44^. We indeed identified variants in several regulators of mitochondria physiology, such as *larp*, *milton*, *Catalase*, together with *MCU* and *MICU1*, two components of mtCU, the mitochondria Calcium Uniporter. This suggests a sensitivity of cardiac aging to variations in mitochondria physiology. Interestingly, our GWAS did not revealed variant in genes encoding core components of mitochondria activity - such as ETC (electron transport chain) components - associated with natural variation of cardiac aging. Not surprising, this observation suggests a strong selection pressure on alleles for these genes, that may have been subjected to selection purification. Overall, our observations demonstrate that, in a natural population, the effects of mitochondria dysfunctions on heart function impact the genetic architecture of its aging. By suggesting that the bZIP transcription factor *Pdp1* – and possibly its paralogue *vri*-controls mitochondria physiology in cardiomyocytes throughout the regulation of some key component that regulate mitochondria homeostasis, our results pave the way towards the identification of a regulatory network involved in heart senescence and regulating mitochondria’s activity.

*Pdp1* and *vri* have been shown to be regulators of the core clock gene *clock* in the central nervous system^28,42^, thus playing an essential role in the circadian oscillator in clock neurons. Whether both genes are also involved in setting the circadian clock in the heart is an open question and it is tempting to speculate that *Pdp1* activity in the heart - and its impact on cardiac aging - may be also related to an effect on the circadian clock. Indeed, several data from mice indicate that aberrant regulation of cardiomyocytes circadian clock contribute to the etiology of cardiac dysfunction and diseases^45^. In flies, reinforcing the circadian oscillator by time restricted feeding was shown to improve cardiac function with age, supporting the contribution of the circadian clock to healthy cardiac aging^46^. Several data from Gill et al^46^ suggested however that an hypothetical *Pdp1* contribution to cardiac intrinsic circadian clock may not be linked to its effect on cardiac aging. Indeed, while we observed a marked increase of AI following cardiac *Pdp1* knock down, loss of function alleles for clock genes displayed no (*clk^ar^*) or opposite effects (*cyc*^01^, *per*^01^ and *tim^01^*) at 5 weeks of age. In addition, neither *Pdp1* nor *vri* cardiac expression was found modified upon time restricted feeding^46^ – which seems to exclude their involvement in improved cardiac function upon circadian rhythm reinforcement.

*Pdp1* has been implicated in a plethora of biological processes. By describing yet another function for this bZIP transcription factor, our study put forward the idea of the pleiotropic function of *Pdp1* in processes as diverse as muscle differentiation^25^, circadian clock regulation^28^, neoplastic growth^47^, organismal growth control^27^ and cardiac functional senescence (this study). Pdp1 is highly homologous to the PAR subfamily of mammalian bZIP transcription factors HLF, DBP and VBP/TEF. Our results therefore may highlight possible processes and diseases affected by these TFs in humans.

## Materials and methods

### Fly strains and husbandry

Fly stocks from the DGRP collection were obtained from Bloomington Drosophila Stock Centre. Flies were reared on standard corn flour medium at 25°C, 50% relative humidity and 12h light/dark cycles. Newly emerged adults (less than 24h) were collected and reared for 7 days or 28 days. 12 flies per DGRP line where dissected and used for high-speed movie capture. The GeneSwitch experiments were done using R94C02::tdTomato (attP2) (hereafter called tdtk)^10^ combined with Hand-GeneSwitch as a stable stock. Driver-line (tdtk; Hand>GeneSwitch) virgins were crossed to UAS-RNAi males (hereafter called KD), UAS-Overexpression (hereafater called OE) males or corresponding isogenic control males. Flies were raised at 25°C on standard fly food. Flies were collected within 24 hours of eclosion under brief CO2 anesthesia and housed in groups of 25. Fly food contains 82.5 mg/ml Yeast, 34 mg/ml Corn meal, 50mg/ml Sucrose, 11,5 mg/ml Agar, 0,5% Methyl 4-hydroxybenzoate (stock solution: 200g/l in ethanol) and RU 232,5 μM (stock solution: 20mg/ml in ethanol) for the RU100 condition (equivalent volume of ethanol was added in the RU0 control condition). Flies were raised at 25°C under a 12hr-12hr light cycle and transferred every two days onto fresh food.

Confocal imaging were realized using Hand-GAL4 driver line (Hand>) virgins, crossed to UAS-RNAi males, Overexpression males or UAS-RNAi control males. ATP quantification and Western Blotting were realized using Mef2-GAL4 driver line (Mef2>) virgins crossed to UAS-RNAi males or UAS-RNAi control males. For these experiments, in order to avoid developmental defects, flies were raised at 18°C on standard fly food throughout development and collected within 24 hours of eclosion under brief CO2 anesthesia and housed in groups of 25. Flies were raised at 25°C for 1 week under a 12hr-12hr light cycle and transferred every two days onto fresh food.

The *Pdp1* deficiency, *Df(3L)BSC631*, was recombined with tdtk as a stable line. All mutant lines were backcrossed into the same isogenic control background. *Df(3L)BSC631*, tdtk and tdtk control virgins were crossed with *MCU^1^* mutant line or isogenic control line. Flies were raised on standard fly food and kept at 18°C. Female progeny were collected and aged to 1 week at 25°C, at which point they were imaged and analyzed.

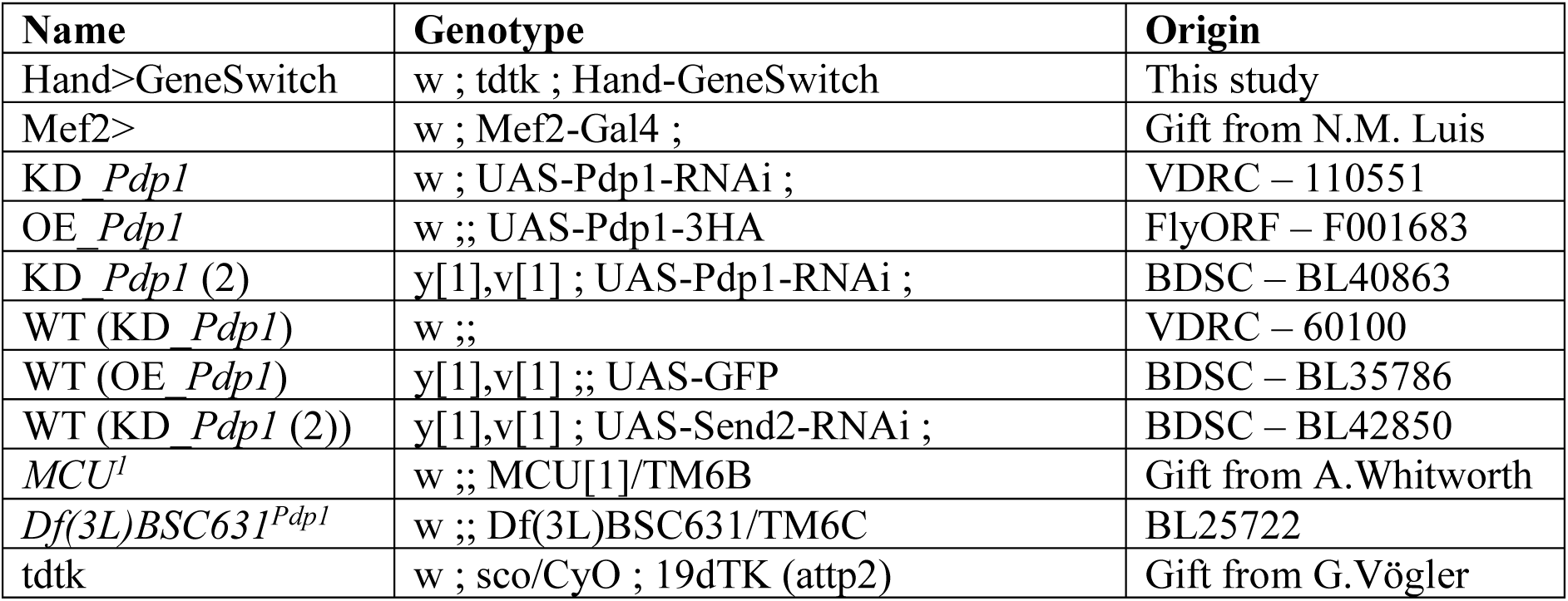

### Semi-intact preparations and SOHA analysis of DGRP strains

Analysis of cardiac function parameters on DGRP flies was achieved on denervated, semi-intact adult female *Drosophila*^19^. 3000 frames of movies were taken at high speed (140fps) with a Hamamatsu ORCAFlash4.0 digital CMOS camera (Hamamatsu Photonics) and using HCI imaging software (Hamamatsu Photonics). These movies were processed using SOHA (http://www.sohasoftware.com/) to measure cardiac parameters.

### Heritability

Heritability is the quantification of the overall phenotypic variation that is attributable to genetic factors. Estimating heritability for complex traits like aging is not straightforward and here we have used a bootstrap approach to obtain heritability due to phenotype aging. For each phenotype, after quality control (see below), from 167 to 175 lines were retained, with 8-12 replicates (individuals) per line at each age. For the DGRP dataset, we cannot track the same fly at age 1 and age 4 because to extract the phenotype, flies are dissected. Hence, we have created bootstrap datasets where within a line, the phenotype for all possible observations at age 1-Week is paired with all possible observations at age 4-Week. For instance, if for line 1, we have 8 observations at age 1 Week and 10 at age 4 Week, then the new bootstrap dataset will have 80 observations, where each age 1 Week observation is paired with each age 4 Week observation. We have used this bootstrap data to obtain the trait difference between age 4 Week and age 1 Week and then calculated the heritability estimates based on the bootstrapped dataset with the trait difference between the two ages. Essentially, the above approach helps us randomly select all possible aging trends from our sample of flies from week 1 to week 4 and determine the heritability of overall aging trend in the DGRP population for each phenotype. We have alternatively also tried to randomly draw 12 observations per line within each age and create a dataset with 167-175 lines (depending on the phenotype) and 12 observations at each age and calculated the heritability estimates based on the trait difference between the two ages. We arrived at approximately similar estimates for heritability with the above two approaches.

### GWAS analyses

Quality Control (QC) was done for each phenotype at each age separately. All phenotype distributions were treated for outliers using standard outlier treatment i.e., removing the observations that lie outside the range [Q1-1.5*IQR; Q3+1.5*IQR], where Q1/Q3 are the first and third quartile of the phenotype values in each line and IQR = Q3-Q1. The removed data were validated by the biologists as technical outliers by looking back at the original experimental results. For GWAS analysis of aging cardiac traits, only those lines that display at least 8 individuals at both 1 and 4 weeks were retained. Depending on the trait, GWAS was performed on 155 to 172 lines (see Table 1).

After quality control, the 8 cardiac traits were compared between 1 week and 4 weeks and except for fractional shortening (FS), there was a strong effect of aging on the phenotype evident from the descriptive plots (box plots) of the phenotype distribution (see Figure 1B). We have also performed quality control on the pooled data from all the lines at each age and removed outliers using a similar method as stated above. In addition, we have conducted a statistical two-sample t-test to test if the phenotype trend varies with age (see Supplemental Figure 1). Almost, all the traits significantly vary with age except fractional shortening. Spearman correlations were also calculated based on the difference between the average of line means at each age for each cardiac trait (see Figure 2E).

Genotype data linked to the cardiac phenotypes were obtained using the GWAS web tool developed for DGRP by Mackay *et al.*^17^. It comprised four million high-quality SNPs among which 95% were homozygous. Typical quality assurance procedures involving maintaining the genotype call (GC) rate at 90% for each chromosome, removing SNPs with minor allele frequency (MAF)>0.019 (that is in 3 lines out of 155, where approximately 155-172 lines were available for GWAS after QC with line observations available at both ages) were performed to get rid of any poor-quality genotype data.

After the QC checks, single marker GWAS was performed on 1,635,932 SNPs from the DGRP data using a linear mixed model (LMM). When testing a marker for association with the phenotype, LMM is a standard application in GWAS, where the variables of interest are modeled as fixed, whereas random effects account for nuisance variation and are integrated out^48^. The linear mixed model included the following covariates: Wolbachia infection status, common polymorphic inversions (data provided by the DGRP Consortium), dates on which the flies were dissected, age of the fly, the SNP (one SNP at a time), the interaction between SNP and age as fixed effects and the strain within each age as random effect. The significance of the interaction between SNP and age in the above model determines the impact of SNP on the aging of the phenotype and helps in determining the list of significant markers that have an impact on aging. SNPs were selected after correction of obtained p values for multi-testing using genome wide Bonferroni adjustment method, the threshold for selection at p< 0,05 was 3 10e-8 (∼0.05/1,635,932). Note that GWAS analysis on DGRP data is primarily conducted on phenotype line means, but in this study, we have used LMM on the individual phenotypes using all the replicates within each line thereby increasing the overall power of the study.

QQ plot were generated to display the quantile distribution of observed marker-phenotype association p-values vs. the distribution of expected p-values by chance. The observed inflation was further investigated. Alternative modeling strategies were tested, as well as an evaluation of our interaction model using a simulation-based strategy. This in-depth analysis is presented in the Supplemental Materials and Methods.

### Annotation of variants according to dm6

Variants present in the DGRP population (VCF file downloadable on the DGRP website: www.dgrp2.gnets.ncsu.edu) were annotated according to dm6 using Variant Effect Predictor (VEP, Ensembl, https://www.ensembl.org/Homo_sapiens/Tools/VEP, default parameters). The following pipeline was then applied to prioritize annotations of each variant. Annotation within one (or more) gene exons were first selected. If the variant was not in a gene exon, annotations within an intron and/or 1kb upstream or downstream of a gene were selected. If the variant was not annotated previously, the closest gene (5kb max) was selected upstream or downstream. If no gene was within this reach, the SNP was annotated in the intergenic region.

### Statistical analysis of DGRP cardiac traits

Phenotypic correlation between each trait pair was computed using Spearman correlation. Pearson chi-squared test was applied to test if the genomic location of variants associated with cardiac traits is biased toward any particular genomic region when compared against the whole set of variants in the entire genome of the DGRP population. Quantification of the overlap between the gene sets associated with the different cardiac traits was done using the overlap coefficient/overlap index.

### Human orthologs

DIOPT v9.0 (https://www.flyrnai.org/diopt, default parameters) was used to identify orthologs of fly and human gene lists. Human gene lists were compiled as described previously^18^ for enrichment analysis of Cardiac Disorder and Coronary Artery Diseases.

Enrichment p-values were based on a test following the hypergeometric distribution. Enrichment (FC) were calculated as the ratio between observed and expected successes in samples (based on the observations in the population).

### G profiler and i-cisTarget analysis

Gene Ontology enrichments were analysed using g:Profiler (https://biit.cs.ut.ee/gprofiler/gost) with default parameters (except Organism = Drosophila melanogaster).

Transcription factor enrichment analysis was done on the genes associated with the End Systolic Diameter (ESD) using i-cisTarget (https://gbiomed.kuleuven.be/apps/lcb/i-cisTarget/index.php) with default parameters.

### In vivo imaging of fly hearts

For *in vivo* imaging of beating hearts, flies were mounted as described^10^ and were illuminated with green light (3 mW power). Five seconds movies of the beating heart were acquired at 300 fps and high-speed recordings were then processed using a custom R script^49^. Beating hearts were imaged with a Hamamatsu ORCAFlash4.0 digital CMOS camera (Hamamatsu Photonics) and using HCI imaging software (Hamamatsu Photonics).

Each experiment was independently normalized. Each individual was normalized to the mean of their control condition, then plotted altogether. Aging trends were modelized using a linear model and compared with an ANOVA test. For single age analysis, a Wilcoxon test was applied between conditions.

### ATP quantification and climbing test

Thoraces of 1 week old female flies were dissected, head and appendices removed. Samples were manually lysed in ATP buffer pH7.8 (100mM Tris, Invitrogen ref:19504020; 4mM EDTA; 6M Guanidine-HCL, Merck ref: G-3272) with a pestle, then snap-frozen in liquid nitrogen before boiling for 5min and centrifuged 13000RPM. Following steps were performed on ice and in 96 wells plates. For each biological sample, quantification of total proteins and of ATP level was performed in parallel. Proteins were quantified using Bradford test (Bio RAD protein assay, ref: 50000). ATP levels were measured using the ATP determination assay (Invitrogen, ref: A22066) according to manufacturer instructions, on VICTOR Nivo plate reader (Perkin Elmer). Data were normalized to protein content. 8 to 10 thoraces were used for each biological replicate (5 for controls and 6 for KD *Pdp1*). 2 independent experiments were performed. Data from Mef2> KD *Pdp1* were normalized to controls Mef2> in each independent experiment. Unpaired Welch’s t test was applied between control and test conditions. Female flies of the same control and test conditions were used for the climbing test video.

### Western blotting

Thorax samples from 10 female flies (1 week old) were dissected, snap-frozen and stored at −80°C until Western Blotting. Lysis was performed in RIPA buffer (Sigma, ref: R0278) with Protease/Phosphatase inhibitor cocktail (Halt™ Protease and Phosphatase Inhibitor Cocktail, ThermoFisher Scientific, ref: 78440). Protein quantification was done using Bradford test (Bio RAD protein assay, ref: 50000). 10ug of proteins were loaded/lane/sample.

Proteins were resolved by SDS-PAGE, using 4-12% gradient gels (NuPage Novex Gel, InVitrogen #NP0335), protein transfert was performed with iBlot2 (InVitrogen, # IB21001) according to manufacturer. Membranes were incubated 2hours in blocking solution (PBS 1x, tween 0,1%, fat-free milk 5%). Primary antibodies used were: mouse anti-ATP5A1 at 1:1000 dilution (ThermoFisher Scientific, ref:43-9800) and mouse anti-Beta actin at 1:1000 dilution (Merck, ref: A5316) in blocking solution O/N at 4°C. Membranes were washed in PBS tween 0,1% and incubated with secondary antibodies in this buffer for 2 hours at RT. Goat anti-mouse HRP conjugated secondary antibody was used at 1:2500 dilution (Thermo Fisher Scientific, ref:G21040). Anti Beta-actin staining was performed on the same membrane after stripping of the ATP5A1 antibody (Restore™ PLUS Western blot stripping buffer, Invitrogen, ref: 46430). Chemiluminescence was observed using the ECL plus detection substrate kit (Thermo Fisher Scientific #32134). Images were generated using Fiji62 (ImageJ version 2.0.0-RC-69/1.52n). Mef2> controls and Mef2> KD *Pdp1* were tested. The experiment was performed 3 times on independent samples.

### Immunochemistry

1 week old female flies were immobilized with stainless steel pins (Fine Science Tools, #26002-10). Flies were then dissected in 1X-PBS (Gibco ref:10010-15) to remove the ventral part of the abdomen, gut, ovaries and expose the heart. For antibody staining, individuals were fixed for 20 min in 1×PBS and 4% paraformaldehyde at room temperature, washed with PBT (1×PBS, 0.3% TritonX-100 (Merck, ref:X100), and incubated for 2 hours with saturation medium, PBT with 10% BSA (Euromedex, ref:04-100-810-C). Primary antibody (mouse anti-ATP5A1 at 1:250 dilution; ThermoFisher Scientific, ref:43-9800) was incubated overnight at 4°C in PBT + 10% BSA. After 4 washes (15’ each) in PBT, samples were incubated 2 hours at RT with Goat anti-mouse Alexa 488 antibody (Life technology, ref:A21042, 1:400 dilution) in PBT. Phalloïdine-555 (Invitrogen, ref:A34055, 1:1000 dilution) was added during the last 20’. Samples were then washed 4 times in PBT before mounting in VectaShield (Vector Laboratories, H-1900).

### Confocal imaging

Observations and images acquisitions were carried out using a LSM 780 Zeiss confocal microscope. Fluorescence quantification was achieved for each heart on 3-4 cardiomyocytes of segment A2 and A3, on Z-stack where the pair of ostia is visible. A square of 200px by 200px for each quantification was used to measure the average fluorescence intensity by measuring for each pixel the grey intensity using <u>Fiji62 (ImageJ version 2.0.0-RC-69/1.52n)</u>. One-way anova with Dunnett’s correction for multiple testing was applied to calculate statistical difference between control and *Pdp1* KD or OE conditions.

### Statistical analysis and Figures design

Statistical analyses and graphical representations were performed with R-Studio/R (RStudio: http://www.rstudio.com/, PBC, Boston, MA; R project version 3.5.1, RRID:SCR_001905) and Graphpad Prism V10 (© 2024 GraphPad Software). The statistical test used for each experiment and the p-values are indicated in the corresponding Figure and Figure legends. Fiji (ImageJ version 2.0.0-RC-69/1.52n) was used for Image treatment from microscopy and Western Blotting. Affinity Designer v.1.10.8 (Serif, Europe) and PowerPoint (Microsoft®) were used for Figures elaboration. We used FlyBase (release FB2025_02) for genes and Drosophila stocks information requests.

## Data availability statement

GWAS data generated in this study are provided in the Supplemental Table 1 (Table of variants with MAF>5% identified on GWAS on the aging of 8 cardiac phenotypes). The analysis code used for GWAS are accessible on Zenodo at https://zenodo.org/records/18911553. List of human orthologues of identified with DIOPT is provided in Supplemental Table 2 (Fly cardiac aging GWAS genes and corresponding human orthologs identified with DIOPT). Raw data of cardiac imaging are provided in Supplemental Datasets 1 (SOHA analysis) and 2 (*Pdp1* functional validation using tdtk method). The authors affirm that all data necessary for confirming the conclusions of the article are present within the article, main figures, tables and supplemental files and figures.

## Acknowledgements

The project leading to this manuscript has received funding from Excellence Initiative of Aix-Marseille University – A*MIDEX (A*Midex International) to L.P., A*MIDEX PEP (AMX-21-PEP-020) to N.A., and ANR (grant n°ANR-22-CE17-0051-03) to N.A.. We thank the Bloomington Drosophila Stock Center (BDSC), the Vienna Drosophila Research Center (VDRC). We also thank A. Whitworth, G.Vögler and N.M. Luis for fly stocks. Centre de Calcul Intensif d’Aix-Marseille is acknowledged for granting access to its high-performance computing resources.

We thank Frederic Gallardo and Tahagan Titus for their technical assistance in fly food preparation. and E. Castellani from the IBDM Imaging Facility and France Bio-Imaging infrastructure - ANR-10 INBS-04-01.

## Author contributions

Katell Audouin: Investigation, Formal analysis, visualization, Writing – editing. Saswati Saha: Data curation, Formal analysis, Visualization, Methodology, Writing – original draft. Laurence Röder: Investigation, Writing – editing. Sallouha Krifa: ressources. Nathalie Arquier: Investigation, Formal analysis, Visualisation, Methodology, Funding acquisition, ressources, Writing – editing. Laurent Perrin: Conceptualization, Formal analysis, Supervision, Funding acquisition, Investigation, Methodology, Writing – original draft, Project administration.

## Conflict of interest

The authors declare no conflict of interest.

## Supplementary figures

**Supplemental Figure 1 (related to Figure 2).** Genome-Wide associations studies (GWAS) for aging of cardiac traits. (A) Reaction norm plot of each DGRP strain at 1 and 4 weeks for ESD, CO, SI and HP. The mean phenotypic values for each line at each age are connected by a line, allowing to monitor the trend differences among DGRP line. (B) Manhattan plot showing the results of GWAS performed on aging phenotypes (ESD, CO, SI and HP). The genome wide significance threshold (3e-8) is indicated by a black line.

**Supplemental Figure 2 (related to Figure 2).** Bar plot of trait evolution during aging for each DGRP strain individually

**Supplemental Figure 3.** QQ of p-values from GWAS on individual phenotypes. Observed SNP-associated p-values were plotted against p-values expected under the null hypothesis. (see Supplemental Materials and Methods for in depth analysis of QQ-plot inflation).

**Supplemental Figure 4 (related to Figure 3).** Heart-specific conditional knock-down (KD) or overexpression (OE) of *Pdp1* modulate cardiac senescence. Cardiac performance was analyzed on intact anesthetized females flies at indicated ages. Number of individuals examined are noted on one plot in color. Individual values were normalized to the mean of the control condition at 1 week. Linear model applied to calculate statistical difference between aging trends of control and condition. (A) Effect of *Pdp1* RNAi mediated gene knockdown (KD). w/w; tdtk/UAS>dsRNA(Pdp1)^VDRC-KK^; Hand>GeneSwitch-Gal4 /+ flies fed (KD-Pdp1) or not (Ctl) with 100µg/ml of RU486 from adult eclosion onward were imaged at 1 week, 4 weeks and 7 weeks. (B) Effect of *Pdp1* over expression (OE). w/w; tdtk/UAS>*Pdp1*; Hand>GeneSwitch-Gal4 /+ flies fed (OE-Pdp1) or not (Ctl) with 100µg/ml of RU486.

**Supplemental Figure 5 (related to Figure 3).** Heart-specific conditional knock-down with an additional RNAi construct (KD-*Pdp1* (2)) modulate cardiac senescence. Cardiac performance was analyzed on intact anesthetized females flies at indicated ages. n= number of individuals examined. Number of individuals examined are noted on one plot in color. Individual values were normalized to the mean of the control condition at 1 week. Linear model applied to calculate statistical difference between aging trends of control and condition. w/w; tdtk/UAS>dsRNA(Pdp1)^BDSC-TRiP^; Hand>GeneSwitch-Gal4 /+ flies fed (KD-Pdp1) or not (Ctl) with 100µg/ml of RU486 from adult eclosion onward were imaged at 1 week, 4 weeks and 7 weeks.

**Supplemental Figure 6 (related to Figure 4).** (A-E) Cardiac performance traits analyzed on adult female flies at 1 week. <u>Control</u>: w/w; tdtk/+; *MCU^1^*: w/w;tdtk/+;*MCU^1^*/+; <u>Df(3L)*Pdp1*</u>: w/w;tdtk/+;Df(3L)BSC631^Pdp1^/+ and MCU^1^/ Df(3L)*Pdp1*: w/w;tdtk/+; MCU^1^/Df(3L)BSC631^Pdp1^. Individual values were normalized to the mean of the control condition. Wilcoxon test was applied to calculate statistical differences between conditions. (A) (corresponding to Figure 4D) Complete uncuted membrane of western blot from thorax lysates (<u>control</u>: Mef2>RNAi-control; <u>KD-*Pdp1*</u>: w/w; Mef2-Gal4/ UAS>dsRNA(Pdp1)^VDRC-KK^). Ponceau red, (visualization of total protein content, left panel), ATP5A1 detection (middle) and Beta-actin detection (right) are shown. (G-I) Confocal imaging of hearts stained for ATP5A1 (green) and filamentous actin (F-actin, red) of control (G: w/w;Hand-GAL4/+); heart specific *Pdp1* knock down (H: w/w;Hand-GAL4/ UAS>dsRNA(Pdp1)^VDRC-KK^) and heart specific *Pdp1* overexpression (I: w/w;Hand-GAL4/+;UAS-Pdp1 /+). Individuals were kept at 18°C throughout development, shifted at 25 °C at adulthood and imaged at 1 week. Longitudinal views of the heart corresponding to Figure 4E-G are shown. VLM=Ventral Longitudinal Muscles, *=Ostia. Scale 10µm. * (p<0,05) / ** (p<0,01) / *** (p<0,001) / **** (p<0,0001)

## Supplementary tables and files

**Supplemental Table 1:** Variant with MAF>5% and their annotations identified by GWAS for senescence of cardiac performance traits at p: 0,05 after genome wide Bonferroni correction (3 10-8 cutoff).

**Supplemental Table 2:** Fly cardiac aging GWAS genes and corresponding human orthologs identified with DIOPT (https://www.flyrnai.org/diopt)

**Supplemental Dataset 1**: Raw data of individual values for cardiac traits in the DGRP. Phenotypic values were determined from high-speed video recording on dissected flies and movie analysis performed using SOHA.

**Supplemental Dataset 2**: Raw and normalized data of individual values for cardiac traits using the high throughput method on intact anesthetized flies. Phenotypic values were determined from high-speed video recording on intact flies expressing tdtk transgene and movie analyzed using tdtk analysis script by Vogler 2022^49^.

**Supplemental Video 1**: Mef2> controls and Mef2>KD *Pdp1,* one week-old female, were compared for their capacity to climb on the vial wall after simultaneous taping at RT.

**Supplemental Materials and Methods**: Statistical analysis of observed QQ-plot inflation. Alternative modeling strategies were tested, as well as an evaluation of our interaction model using a simulation-based strategy.

## Notes

### Competing Interest Statement

The authors have declared no competing interest.

### Summary of Updates

This revision includes: new statistical analysis for the GWAS data, new experiments regarding the Pdp1 characterization.

