## Supplemental Material and Methods for "Natural variations of cardiac performance in *Drosophila* identify a central function for *Pdp1*/*dHLF* in cardiac aging"

### Supplemental Materials and Methods

#### QQ-plots analyses

The following model was fitted in GWAS:

$$y_{ijk} = \beta_0 + \beta_1 date_{ijk} + \beta_2 (snp \times age)_{ik} + \beta_3 snp_i + \beta_4 age_k + \sum_m \gamma_m Inv_{ijm} + u_i + \epsilon_{ijk}$$

Where

- $y_{ijk} = phenotype\ value\ for\ strain\ i, age\ k\ and\ observation\ j\ within\ strain\ i.$
- $u_i \sim N(0, \sigma^2_{strain}) = random\ effect\ due\ to\ strain(intercept)$
- $\epsilon_{ijk} \sim N(0, \sigma^2) = residual\ variance$
- $\beta_1$ was the effect due the date on which the flies are dissected. Each strain was dissected at different dates and the dates were different from age 1 to age 4.
- $\beta_2\ is\ effect\ due\ to\ the\ interaction\ between\ snp\ and\ age.$
- $\beta_3\ is\ the\ effect\ due\ to\ snp.$
- $\beta_4\ is\ the\ effect\ due\ to\ age.$
- $\gamma_m\ is\ the\ effect\ due\ to\ the\ inversion$

We focused on testing the significance of $\beta_2$ in our GWAS. The p-values were obtained from the interaction GWAS conducted on 1,635,932 SNPs. To control for false positives, we applied a stringent Bonferroni correction by dividing 0.05 by 1,635,932, resulting in a genome-wide significance threshold of $3 \times 10^{-8}$.

The QQ plot of the GWAS p-values across all phenotypes showed some inflation. However, it is well known that a certain degree of inflation is commonly observed in interaction GWAS analyses[1,2], particularly due to reduced power and model complexity.

To investigate this further, we extracted phenotype data for *Heartperiod Median* and performed sensitivity analyses. Specifically, we selected SNPs from chromosome 2L of the *Drosophila Genetic Reference Panel* genome dataset and repeated the GWAS sensitivity analysis restricted to these variants.

We observed that the inflation persisted at the chromosome level. Because repeating the sensitivity analysis genome-wide would have been computationally intensive, we limited these additional analyses to chromosome 2L, which represents approximately 20% of the genome in terms of number of SNPs (356,273 SNPs). The results for chromosome 2L are presented here.

We first examined whether the specification of the standard errors—particularly the definitions of the *u*, *v*, and *ε* components—might have contributed to the observed inflation. To assess this, we implemented three alternative modelling strategies:

- Lineaer Mixed Models (LMM) fitted on line means
- LMM incorporating a genetic relationship matrix (GRM)
- LMM using robust sandwich standard errors

##### *LMM on Line Means*

We repeated the GWAS using the same model structure as in the primary analysis, but instead of fitting it at the individual level, we first computed phenotype means across all individuals within each line and age group.

We then fitted the following model:

$$y_{ik} = \beta_0 + \beta_1 date_{ik} + \beta_2 (snp \times age)_{ik} + \beta_3 snp_i + \beta_4 age_k + \sum_m \gamma_m Inv_{im} + u_i + \epsilon_{ik}$$

The results were essentially identical: the p-values from the line-mean GWAS closely matched those from the individual-level GWAS using the original model. QQ plots from both the analysis are shown below (associated Figure 1&2 and plot of p value concordance Figure 3).

##### *LMM incorporating a genetic relationship matrix (GRM)*

In the original analyses above, the kinship matrix K was assumed to be an identity matrix. One possible source of inflation could therefore be the omission of relatedness (i.e., the genomic relationship/kinship matrix) from the GWAS model.

To test this, we repeated the GWAS on chromosome 2L while incorporating the kinship matrix i.e. assumed in the above model,

- $u_i \sim N(0, \sigma^2_{strain} K) = random\ effect\ due\ to\ strain(intercept)$

Previously, kinship was not included because we did not expect strong population structure effects in the DGRP data, and because inclusion of the GRM substantially increased computation time. However, incorporating the kinship matrix had little impact on the GWAS results.

##### *LMM using robust sandwich standard errors*

Inflation of p-values in interaction GWAS that include genotype-by-environment (G×E) terms can arise from phenotype heteroscedasticity across environmental strata[1]. Standard linear mixed models assume homoscedastic residual variance across environmental levels, an assumption that may not hold in practice.

Because our model includes a genotype × age interaction and phenotypic variability changes across age groups, variance heterogeneity represented a plausible source of model misspecification and the resulting p-value inflation. We therefore repeated the GWAS using the alternative error structures recommended in the cited study[1]. However, the results were largely unchanged, and p-value inflation persisted (see associated Figure. 4).

Because multiple sensitivity analyses and alternative model specifications did not identify any evident source of confounding, we concluded that the observed inflation was unlikely to result from model misspecification. This led us to examine a more fundamental question: whether the assumed null distribution of the interaction test statistic is correctly specified in this context.

In conventional GWAS analyses, interaction test statistics are typically evaluated against a central Student's t distribution (equivalently, under large-sample assumptions, the squared test statistic is assumed to follow a $\chi^2$ distribution). However, recent work[3] describing the "feast and famine" phenomenon in interaction GWAS fitted with linear mixed models demonstrates that, under the null hypothesis (when there is no association between the phenotype and genotype at any level), the interaction test statistic does not necessarily follow a standard central *t* distribution. Instead, it follows a *t*-distribution with mean zero but variance $v$, where $v$ is not constant and depends on the observed phenotype values and the covariates included in the model.

If the true null distribution deviates from the conventional central *t* (or $\chi^2_1$) distribution, then interpreting apparent p-value inflation using the traditional genomic inflation factor ($\lambda$)—which assumes $\chi^2_1$ behaviour under the null—may be inappropriate for interaction GWAS. This issue has also been discussed in literature[4], where the authors show that variability in $\lambda$ can arise from non-independence among test statistics coupled with stochastic bias due to incomplete asymptotic behaviour of the test statistic.

To further evaluate the validity of our interaction model, we adopted a simulation-based strategy motivated by prior methodological work[4,5]. We generated phenotype data under the null hypothesis by enforcing identical aging trajectories for the heart period phenotype from week 1 to week 4 across both minor and major alleles, thereby eliminating any true genotype-by-age interaction effect (see associated Figure 5) while preserving the main effects of age and genotype on the phenotype. We then repeated the GWAS on chromosome 2L using the simulated phenotype and extracted the corresponding interaction test statistics and p-values. This procedure yielded an empirical null distribution for the interaction test statistic under our modelling framework.

We next derived empirical p-values by placing the observed interaction test statistics from the original GWAS (based on the real phenotype data) in the context of the simulated null distribution. Specifically, for each SNP, the empirical p-value was calculated as the proportion of test statistics from the simulated null that were equal to or more extreme than the observed test statistic (see associated Figure 6 for more details). Notably, the ranking of SNPs based on empirical p-values was identical to that obtained from the model-derived p-values in the original analysis, with a Spearman rank correlation of 1.0 between the two sets of p-values.

The fact that SNP ranking is perfectly preserved indicates that the strongest signals observed in the original GWAS are not artifacts of mis-specified model assumptions. Rather, the relative extremeness of the top interaction statistics is maintained even when evaluated against an empirical null tailored to our data structure. Taken together, these results provide strong evidence that the leading interaction hits identified in our GWAS are unlikely to represent false positives.

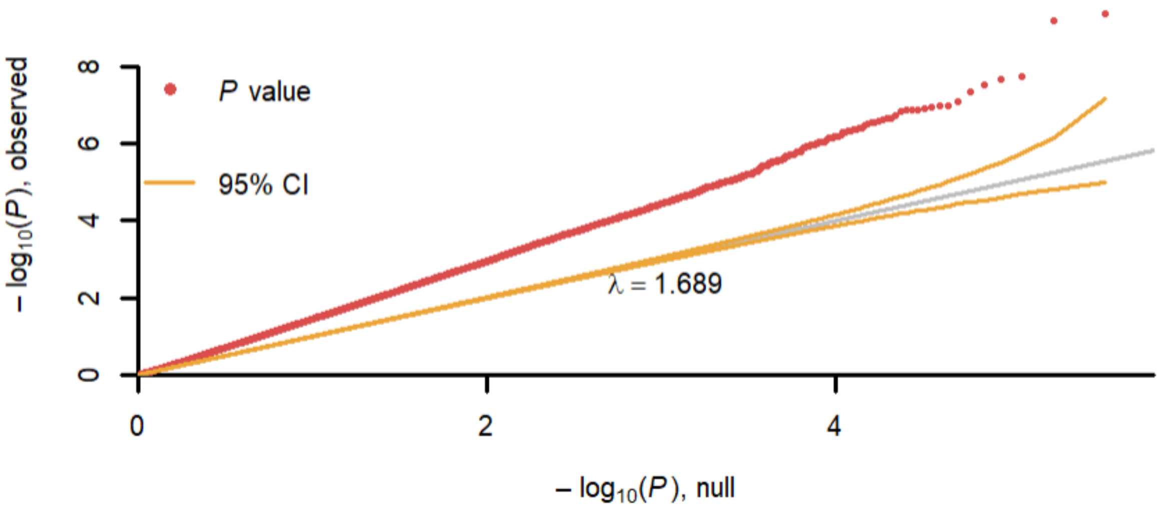


**Figure 1: QQPlot of p-values from individual GWAS (Chromosome 2L - 356273 SNPS)**

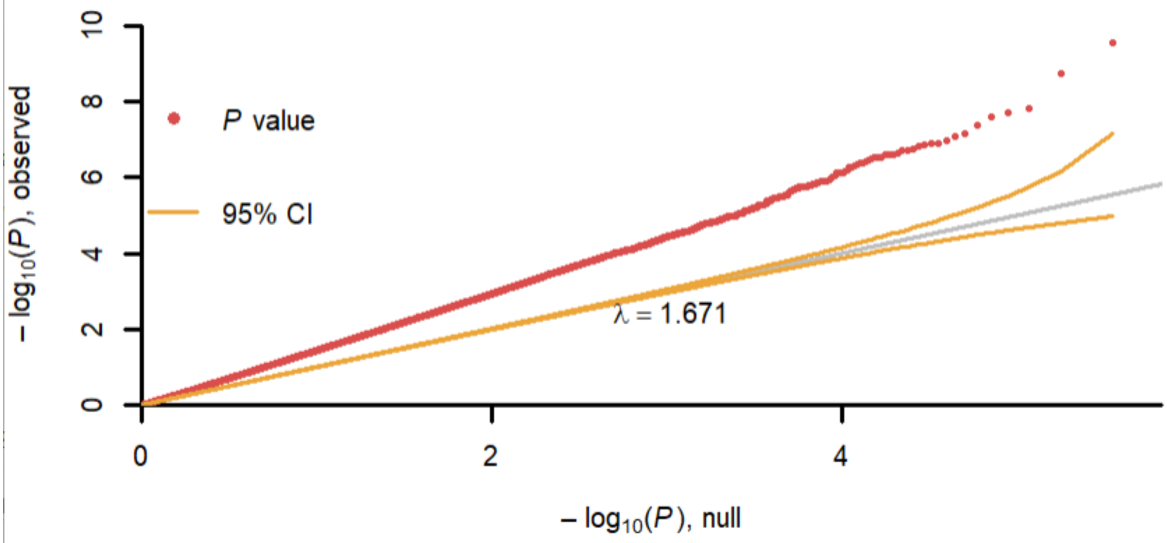


**Figure 2: QQPlot of p-values from line mean GWAS (Chromosome 2L - 356273 SNPS)**

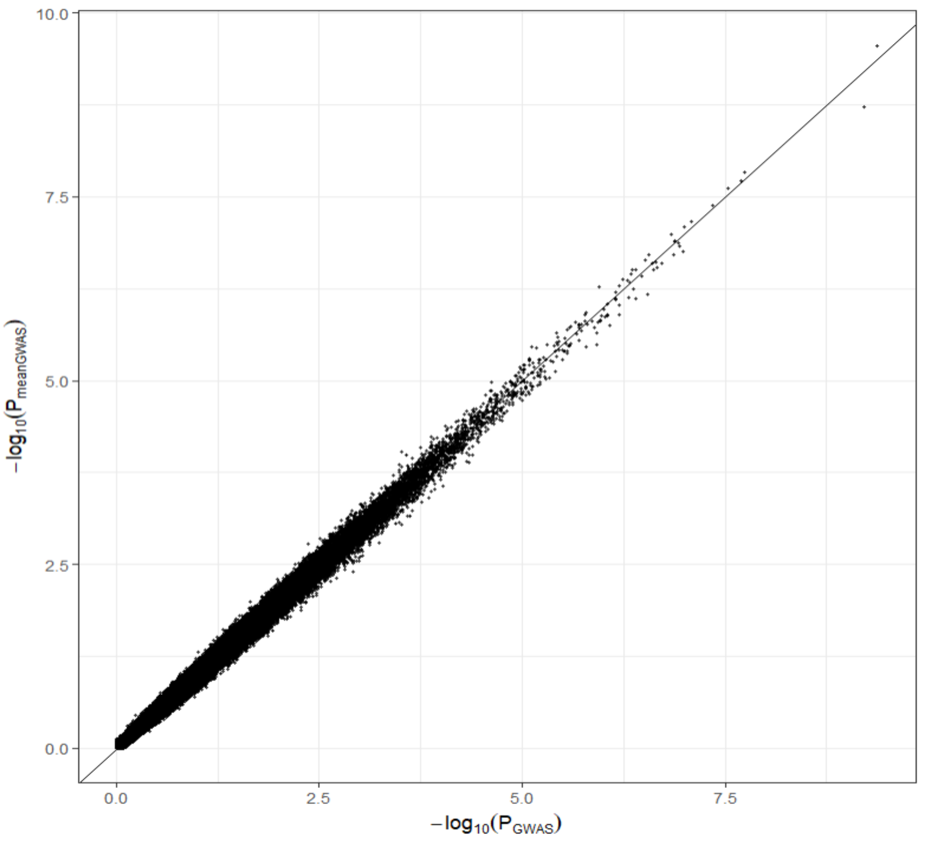


**Figure 3: Concordance between line mean and individual GWAS p-values (Chromosome 2L - 356273 SNPS).**

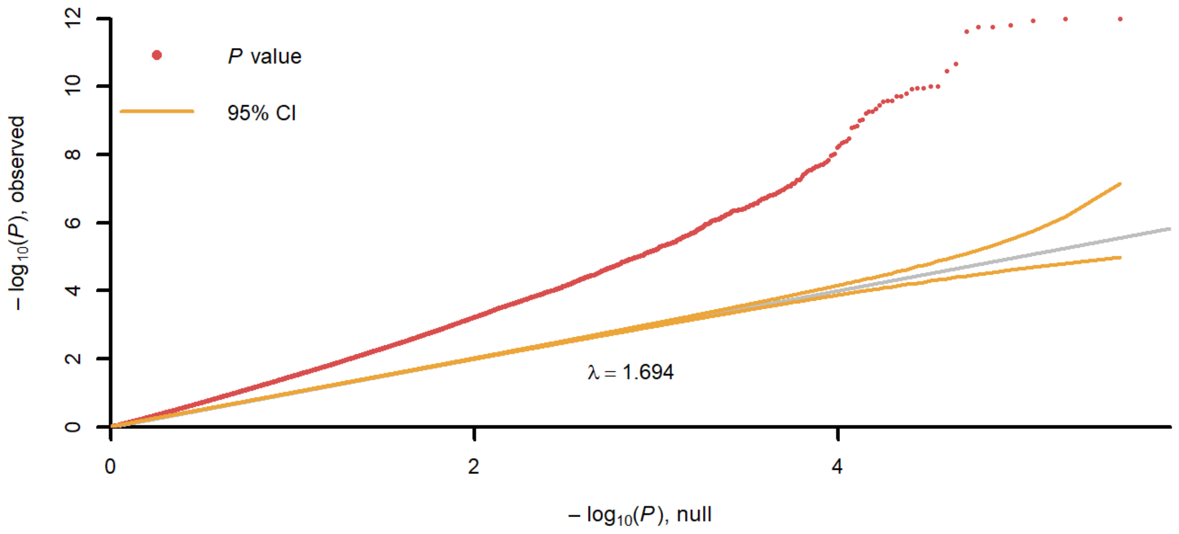


**Figure 4: QQPlot of p-values from alternative error structures for interaction models (Chromosome 2L - 356273 SNPS)**

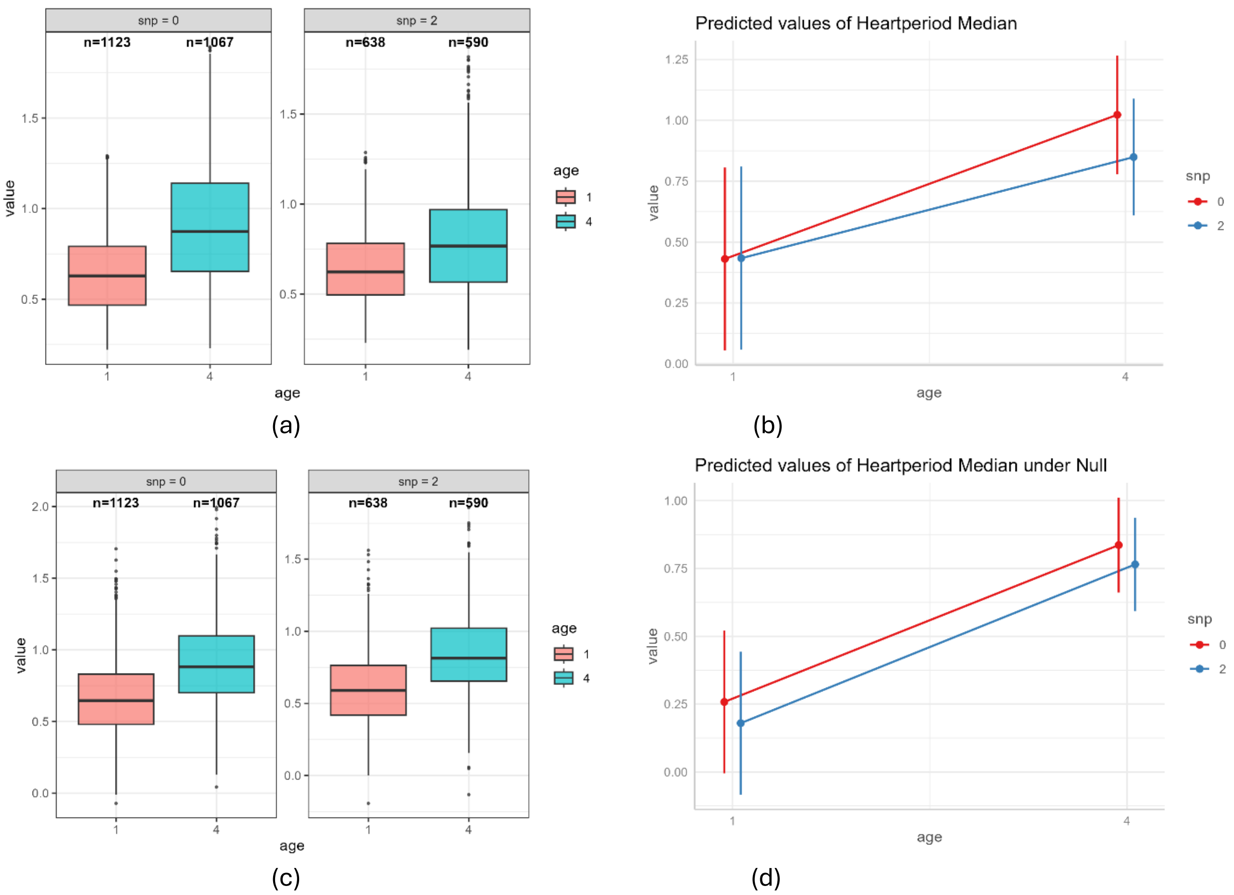


**Figure 5: The simulated null data do not show that the simulated data under null eliminated the genotype-age interaction that is there in the original data. (a)** Distribution of Heartperiod Median at 1 and 4 weeks of age, stratified by genotype at SNP position 15,105,255 on chromosome 2L (SNP = 0 vs. SNP = 2). Sample sizes for each group are indicated above the boxplots. **(b)** Predicted mean Heartperiod Median at 1 and 4 weeks for each genotype group (SNP = 0 and SNP = 2), illustrating the age-related trend and the difference in aging patterns between genotypes. **(c)** Distribution of Simulated Heartperiod Median under Null at 1 and 4 weeks of age, stratified by genotype at SNP position 15,105,255 on chromosome 2L (SNP = 0 vs. SNP = 2). **(b)** Predicted mean Simulated Heartperiod Median under Null at 1 and 4 weeks for each genotype group (SNP = 0 and SNP = 2), illustrating the age-related trend and the difference in aging patterns between genotypes.

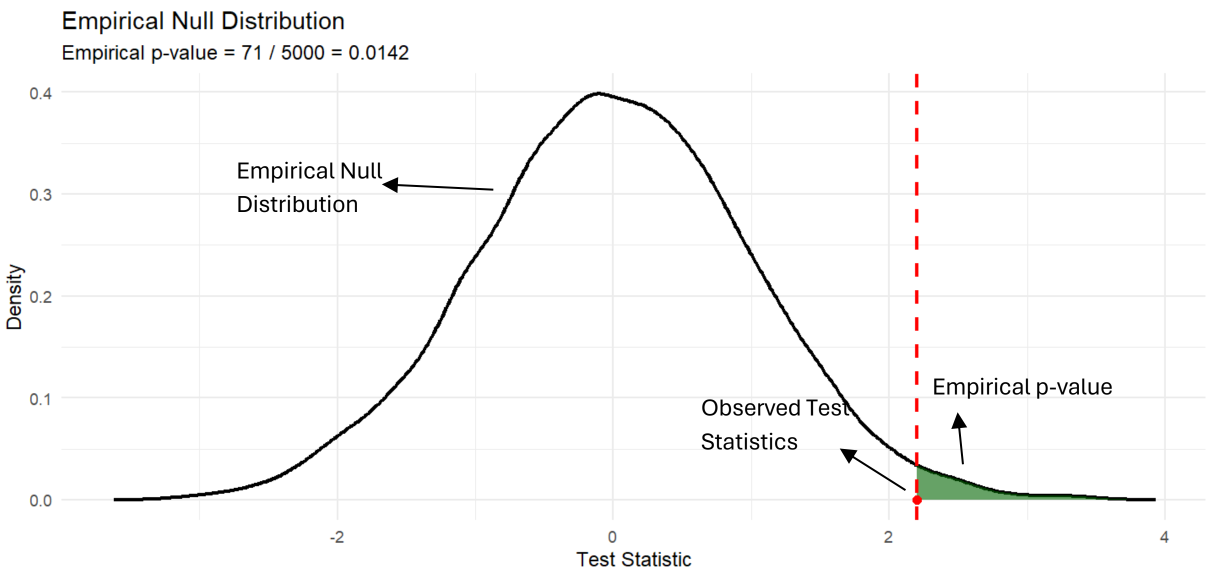


**Figure 6**: The plot shows the empirical null distribution of the test statistic, with the shaded tail representing the proportion of simulation-based null values that are more extreme than the observed statistic—i.e., the empirical p-value.
