## Supplementary figures and images for "Natural variations of cardiac performance in *Drosophila* identify a central function for *Pdp1*/*dHLF* in cardiac aging"

### Supplemental Figure 1

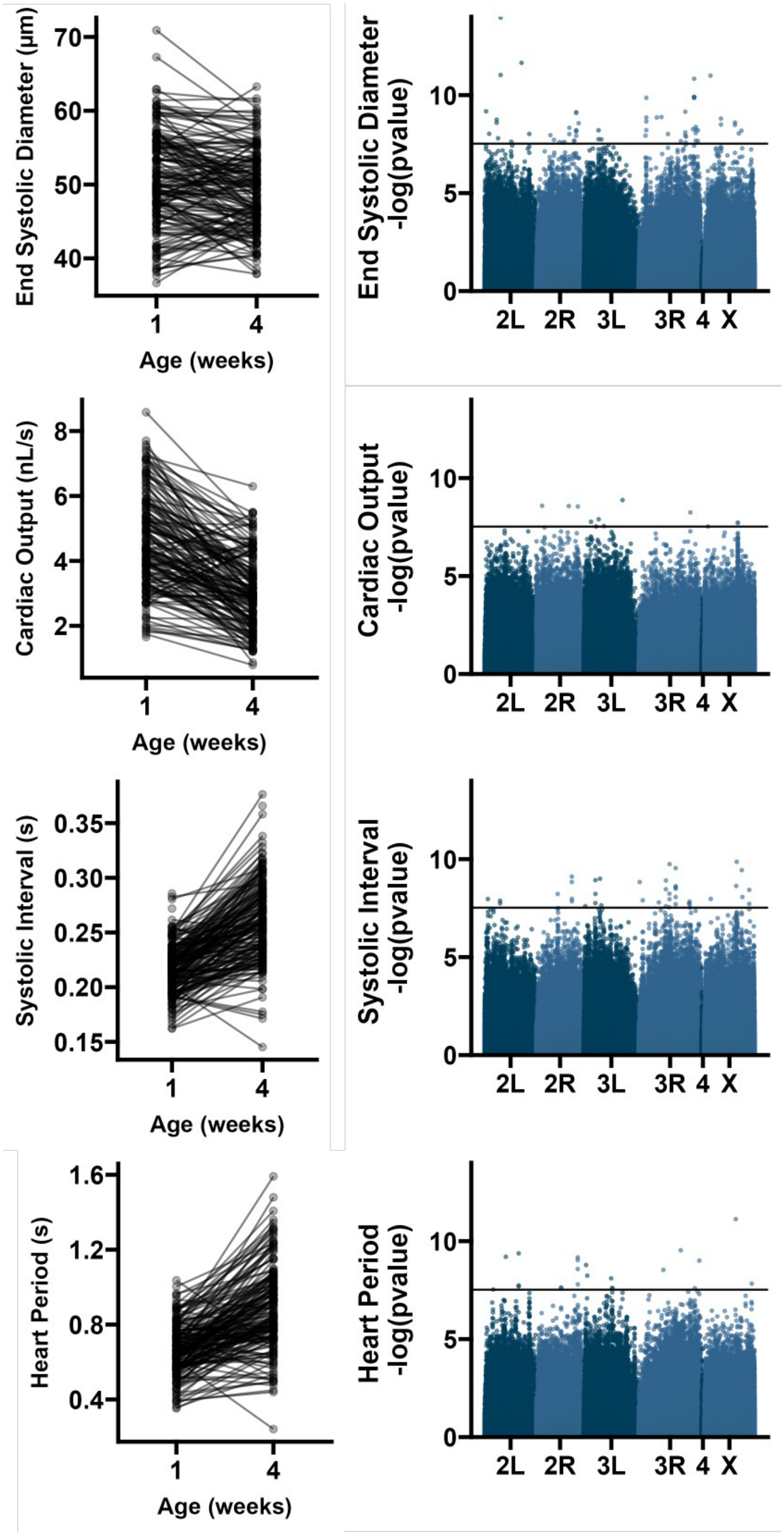


**Supplemental Figure 1**

### Supplemental Figure 2

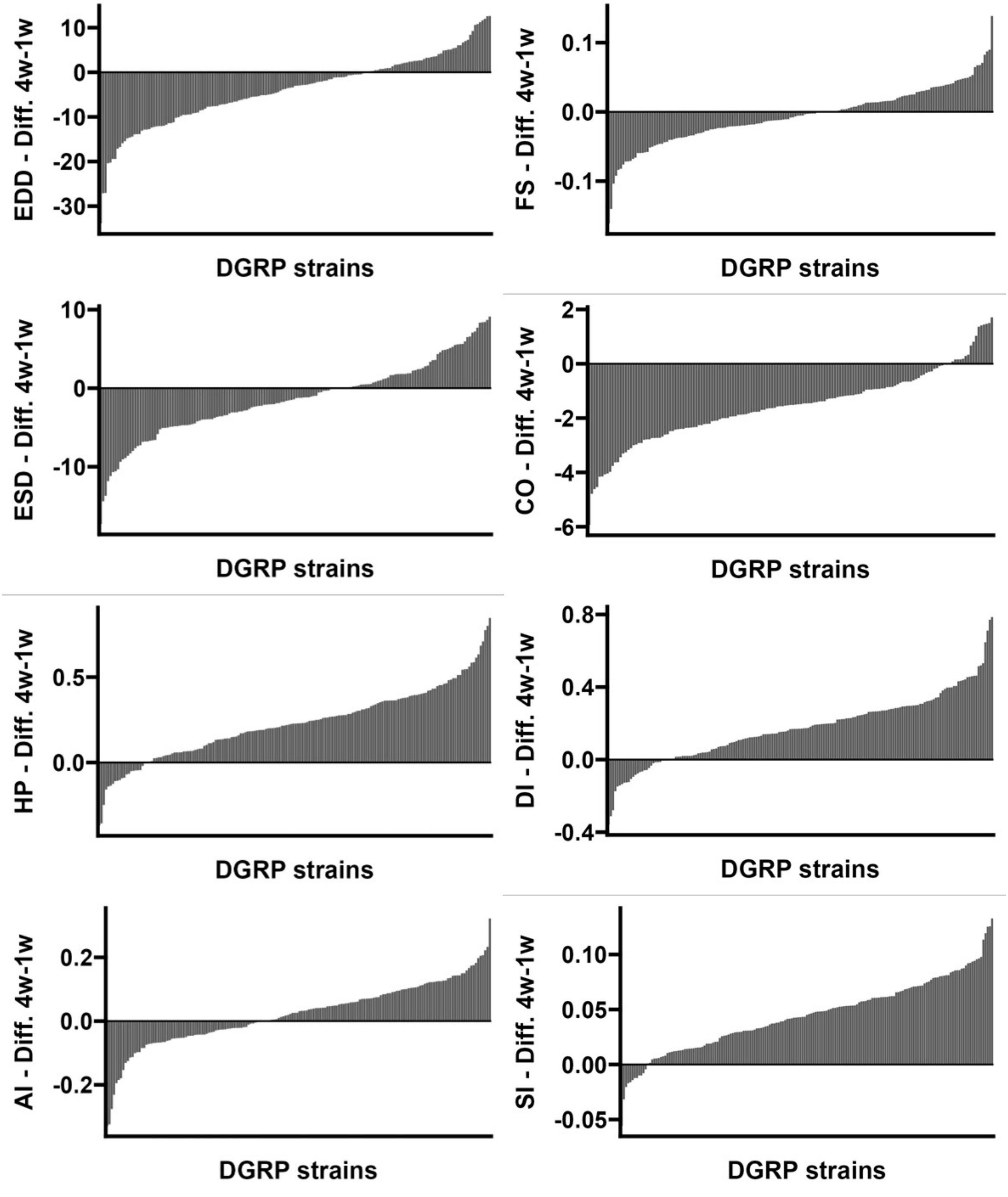


**Supplemental Figure 2**

### Supplemental Figure 3

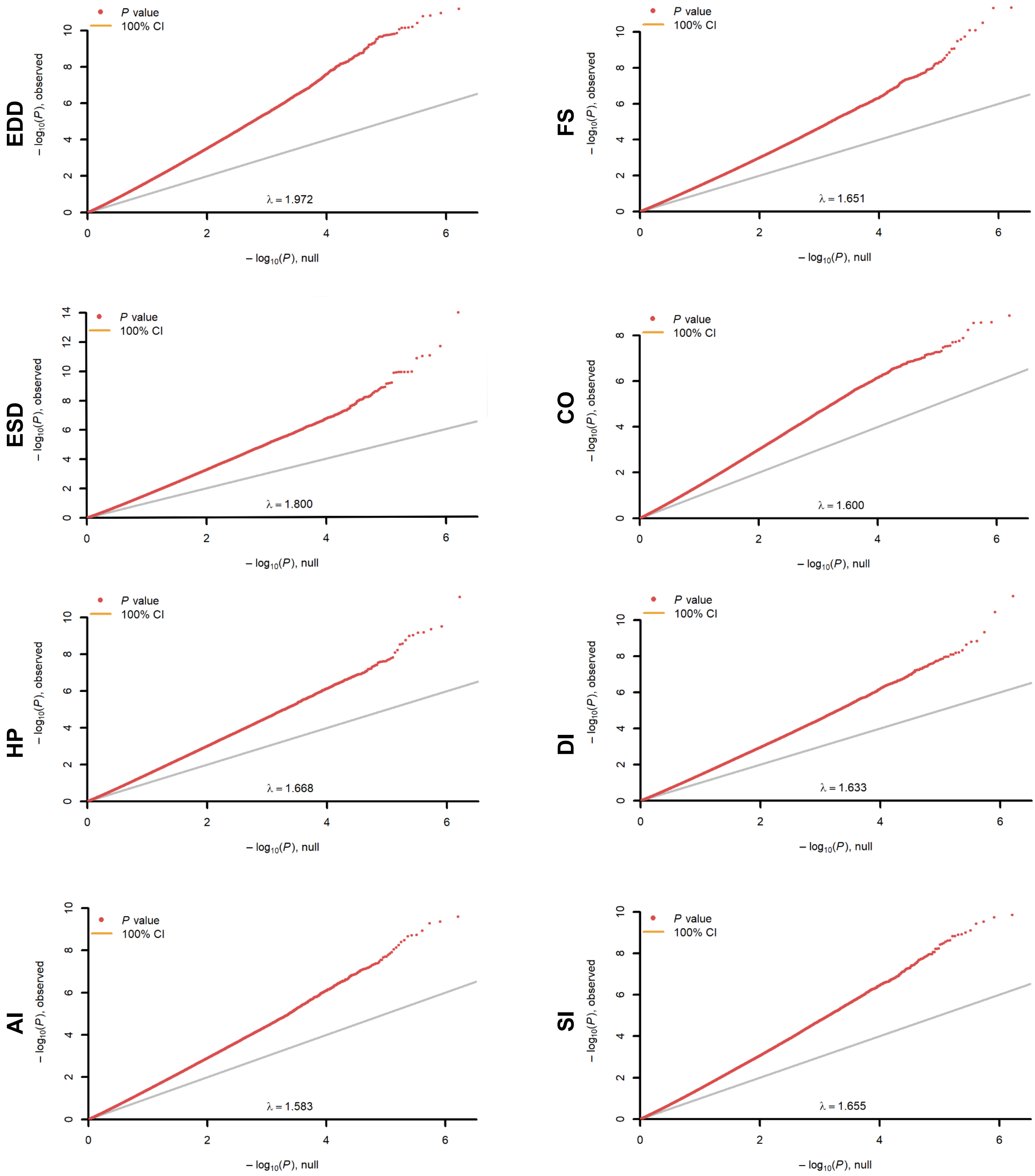


#### Supplemental Figure 3

### Supplemental Figure 4

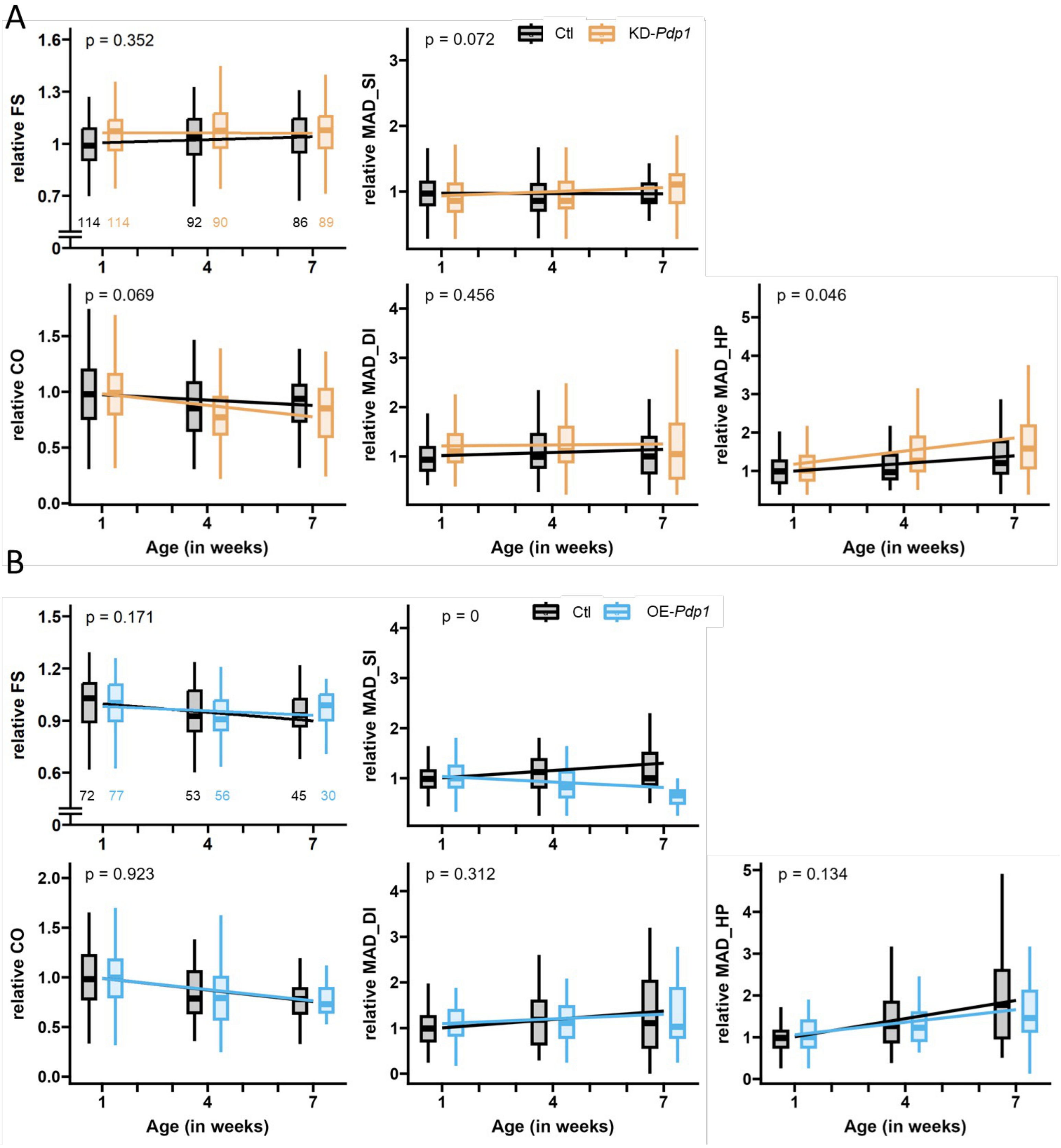


**Supplemental Figure 3**

### Supplemental Figure 5

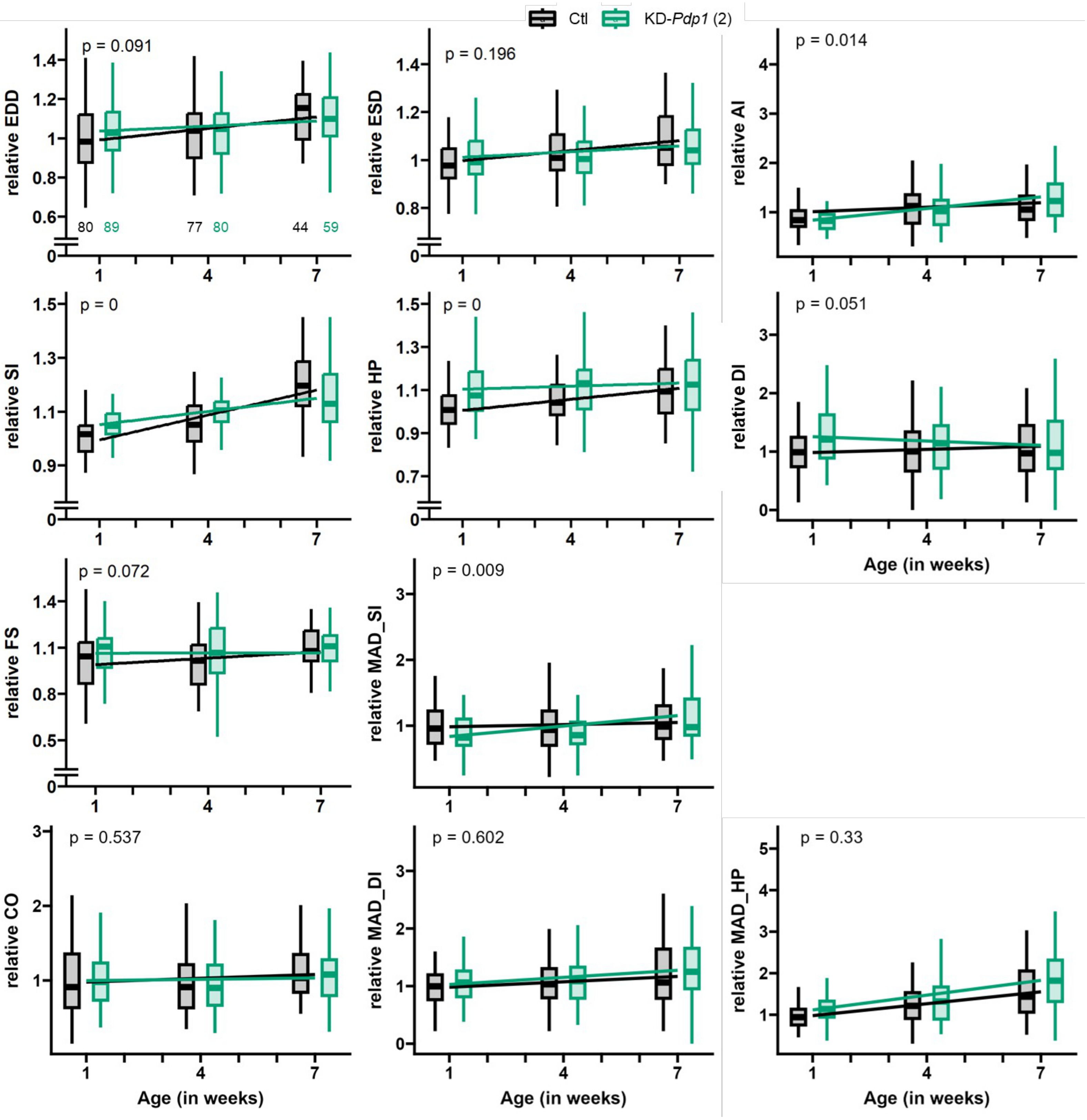


**Supplemental Figure 4**

### Supplemental Figure 6

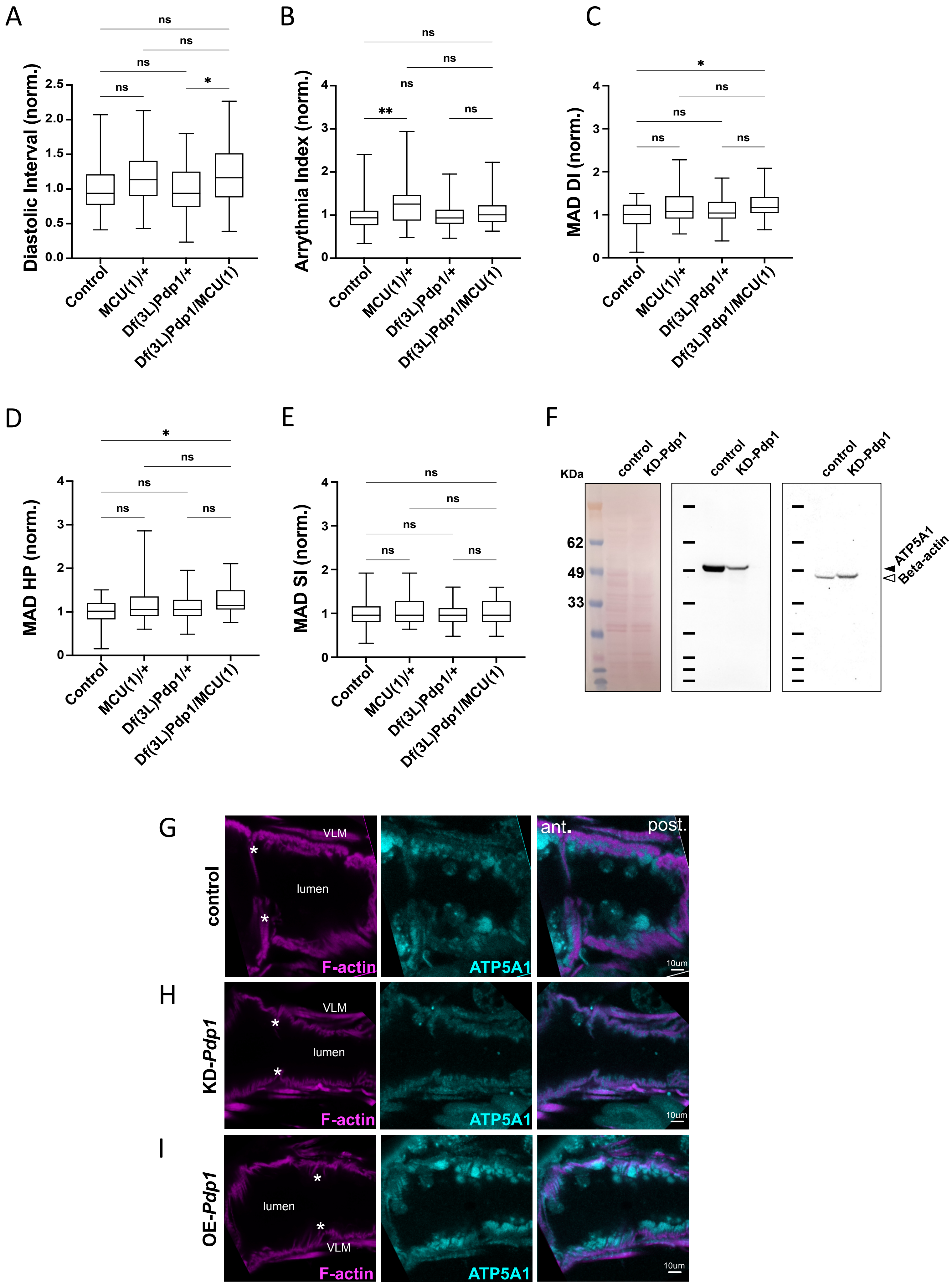


Supplemental Figure 5
